# AptaBLE: A Deep Learning Platform for Aptamer Generation and Analysis

**DOI:** 10.64898/2026.01.06.698056

**Authors:** Sawan Patel, Keith Fraser, Dhanush Gandavadi, Abhisek Dwivedy, Xing Wang, Fred Zhangzhi Peng, Pranam Chatterjee, Sherwood Yao

**Affiliations:** Atom Bioworks, Cary, NC, USA; Department of Biological Sciences, Rensselaer Polytechnic Institute, Troy, NY, USA; Center for Biotechnology and Interdisciplinary Studies, Rensselaer Polytechnic Institute, Troy, NY, USA; Future of Computing Institute, Rensselaer Polytechnic Institute, Troy, NY, USA; Department of Bioengineering, Grainger College of Engineering, University of Illinois at Urbana-Champaign, Urbana, Illinois, USA; Holonayak Micro & Nanotechnology Lab, Grainger College of Engineering, University of Illinois at Urbana-Champaign, Urbana, Illinois, USA; Carl R. Woese Institute for Genomic Biology, University of Illinois at Urbana−Champaign, Urbana, Illinois, USA; Department of Chemistry, University of Illinois at Urbana−Champaign, Urbana, Illinois, USA; Cancer Center at Illinois, University of Illinois at Urbana−Champaign, Urbana, Illinois, USA; Department of Biomedical Engineering, Duke University, Durham, NC, USA; Department of Computer and Information Science, University of Pennsylvania, Philadelphia, PA, USA; Department of Bioengineering, University of Pennsylvania, Philadelphia, PA, USA

## Abstract

Aptamers are single-stranded oligonucleotides that bind molecular targets with high affinity and specificity. However, their discovery remains time-consuming, expensive, and susceptible to experimental biases. Here we present AptaBLE, a deep learning framework for predicting aptamer-protein binding. Additionally, we demonstrate two *de novo* generation methods that produce novel aptamers with desired specificity profiles and K_d_’s as low as 31 nM to-date. AptaBLE represents a significant advance towards therapeutic and diagnostic aptamer development.

---

Among therapeutic modalities, aptamers, single-stranded DNA or RNA oligonucleotides that fold into complex three-dimensional structures, have emerged as promising alternatives to antibodies^1,2^. Still, aptamer discovery remains constrained by the limitations of Systematic Evolution of Ligands by EXponential enrichment (SELEX)^3^. The SELEX process involves iterative rounds of selection, amplification, and enrichment from randomized oligonucleotide libraries containing between 10^14^ and 10^15^ unique sequences^3^. Typical campaigns require 10-15 rounds of iteration over several months and can often be complicated by sequence bias introduced through the necessary per-step polymerase chain reaction (PCR) amplification^4,5^. This is exacerbated by traditional library analysis methods reliant on sequential homology clustering^6,7^.

Such limitations have motivated the development of computational approaches to process aptamer libraries^8,9^, though they provide limited mechanistic insight. Advances in deep learning offer opportunities for aptamer design^10-14^. However, existing methods poorly model long-range intermolecular interactions. Additionally, the scarcity of solved aptamer-protein structures, and the inherent flexibility of nucleic acid polymers^15,16^, creates biases in structure-based approaches^17-21^ which are reflected by historically poor performance on nucleic acid modalities.

To address this, we present **AptaBLE** (**Apta**mer **B**inding **L**anguag**E**; **Figure 1A**), a deep learning framework capable of both predicting the binary interaction of aptamer-protein complexes and generating high-affinity aptamers *de novo*.

**Figure 1.**
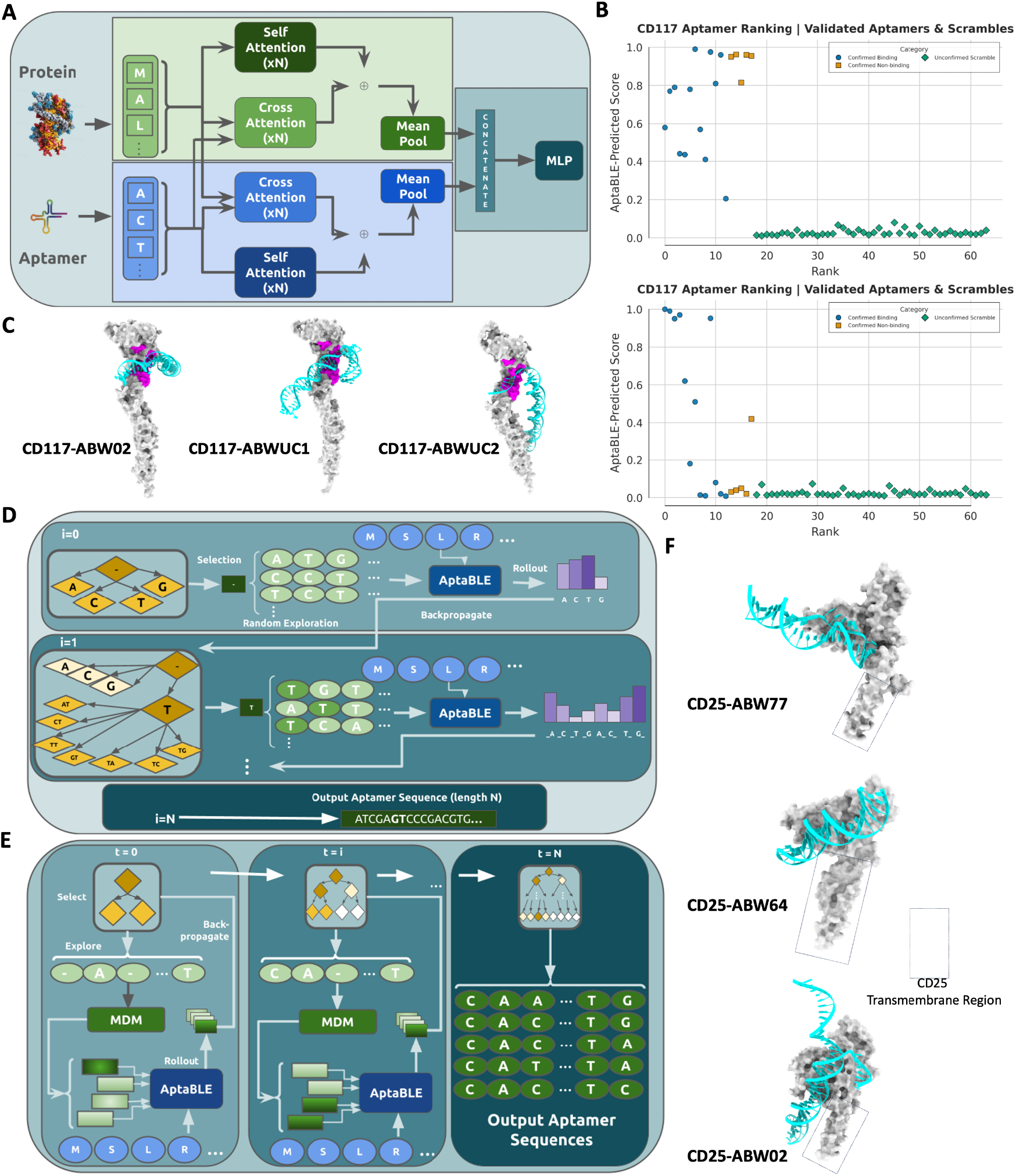
AptaBLE architecture and *in silico* benchmarking. **A)** Protein and nucleic acids are represented by their primary structures and are embedded via pretrained encoders (**left**). The fusion module combines embeddings into a shared representation, then downsized to a score with an MLP (**right**). **B)** Aptamers predicted to bind human CD117 are plotted by AptaBLE score alongside their log-*K*_*d*_, if determined **(Top**). Confirmed binding aptamers (blue) are ordered by their *K*_*d*_. A set of non-binding aptamers are also shown (orange) in addition to 64 scrambled sequences (green). **(Bottom)** After fine-tuning on two binding aptamers (ABW02, ABW04), an approximate ordering by *K*_*d*_ emerges. **C)** Molecular docking comparison between aptamers ABW02, ABWUC1, and ABWUC2 in complex with CD117, with its stem cell factor binding site used to facilitate receptor dimerization highlighted (magenta). **D)** Algorithm schematic for AptaBLE-MCTS (see **Algorithm S1** for details). **E)** Algorithm for AptaBLE-MCTG (see **Algorithm S2** for details) **F)** Molecular docking study of Aptamer 77 in complex with hCD25. Dashed box denotes the transmembrane region of the receptor.

First, we performed a retrospective analysis on a human CD117 (hCD117) aptamer SELEX library^22^. We evaluated model performance on a series of hCD117-selective aptamers with known binding affinities. This diagnostic comprises 13 binding aptamers, 5 validated non-binding aptamers and a series of scrambled sequences. We show that the majority of binding aptamers fall above a 0.4 score, with the least-binding aptamer falling below the 0.4 threshold (**Figure 1B, top**). Notably, all of the validated non-binding aptamers score highly, as well as a small subset of scrambles. We see that there is no observable correlation with experimental *K*_*d*_ ‘s (**Figure S3, left**). Upon fine-tuning AptaBLE with a subset of two hCD117 aptamers, ABW02 and ABW04, the previously high-scoring scrambles sequences are then marked as poor binders (**Figure 1B, bottom**). We also observe a relatively stronger correlation and ordering of the AptaBLE scores for each aptamer according to their experimentally-determined *K* _*d*_’s after fine-tuning (**Figure S3, right**).

The above 12 binding aptamers stem from prior work where we evolved ssDNA sequences binding to hCD117 through nine rounds of SELEX^25^. In an effort to retrospectively process the product library, we used AptaBLE to parse through the population and identify potential high-affinity binders that were missed. We leveraged a combined approach utilizing AptaBLE scores and sequence copy number (**Figure S4A**). While we focus on unclustered aptamers here, we also scored all clustered aptamers and the overall score distributions across clusters (**Figure S4B,C**). Setting the thresholds for AptaBLE score to 0.4 and a minimum copy number of 10, we identified a sub-population of the unclustered aptamers with potential as binders specific to hCD117. We chose four aptamers, according to their predicted minimum free energies in testing buffer conditions and distinct structures to ABW02, an existing high-affinity hCD117 binder^22^, according to a hierarchical clustering analysis of their predicted secondary structures (**Figure S5**). These unclustered aptamers were all found to bind hCD117 (**Table S1**). Indicated as ‘ABWUC’ followed by an arbitrary number, these aptamers showed, in some cases, favorable binding kinetics over previously tested aptamers selected via the traditional library analysis (ABW02-ABW07). ABWUC1 and UC2 achieve sub-50 nM affinity to hCD117. Predicted docking complexes for ABW02, ABWUC1, and ABWUC2 with hCD117 are shown in **Figure 1C**.

Subsequently, we sought to exploit the model’s predictive capability to design aptamers *de novo* for targeting cell surface receptors by adapting existing algorithms^23,24^. Our two methods, named AptaBLE-MCTS (**Figure 1D**) and AptaBLE-MCTG (**Figure 1E**), were evaluated in tandem. We generated and shortlisted five candidate aptamers for targeting the immune receptors TIGIT and CD25 (see Supplementary: *<u>De Novo Generation</u>*). Initial screening of all candidate aptamers was performed at a fixed concentration of 10 μM to assess binding-induced fluorescence shifts relative to untreated controls (**Figure S6**). Among the tested sequences, Aptamers 64 and 77 showed the highest fluorescence shifts, indicating relatively superior target engagement. For further quantitative binding analysis, we selected Aptamers 22 and 27 as anti-TIGIT candidates and Aptamers 64 and 77 as anti-CD25 candidates. Molecular docking complexes for Aptamers 64, 77, and ABW02 with CD25 are provided in **Figure 1F**.

Anti-TIGIT Aptamers 22 and 27 were evaluated at 310 K (37 °C) for 40 minutes over a concentration range of 0.1–20 μM in TIGIT-positive MM1.S cells and TIGIT-negative HEK293T cells (**Figure 2A, S7**). Both aptamers exhibited strong, concentration-dependent binding to MM1.S cells while minimal binding was observed in HEK293T cells. Binding curve analysis yielded dissociation constants (K_d_) of 7.3 μM for Aptamer 22 and 12.55 μM for Aptamer 27.

**Figure 2.**
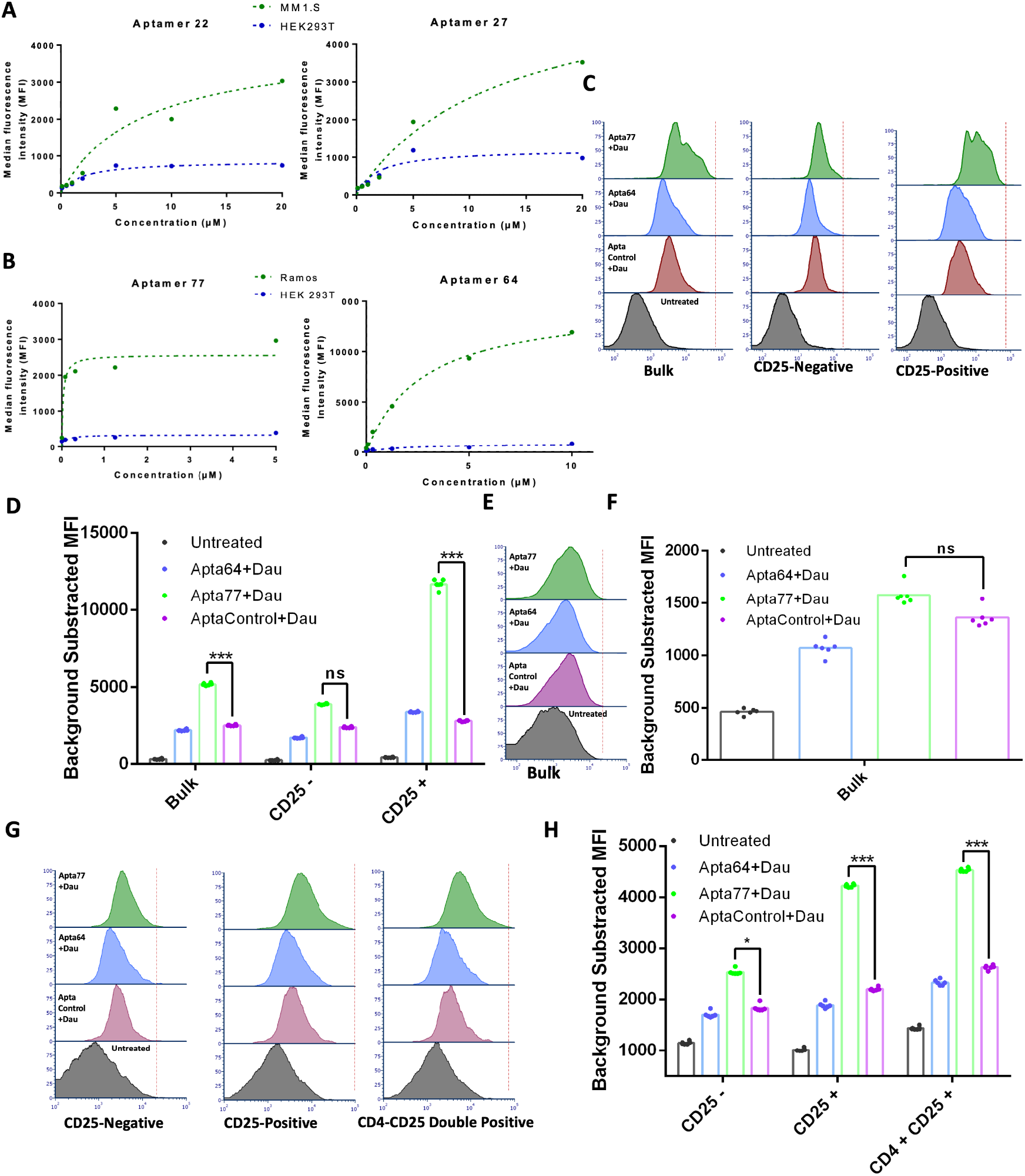
AptaBLE-designed aptamers bind selectively and with low-nanomolar affinity to their target protein *in vitro*. **A)** Concentration-dependent binding of anti-TIGIT aptamers to TIGIT-positive MM1.S cells (green) and TIGIT-negative HEK293T cells (blue) measured by median fluorescence intensity (MFI) following incubation at 310 K for 40 min. Binding curves are shown for Aptamer 22 (left; *K*_*d*_ ∼7.3 μM) and Aptamer 27 (right; *K*_*d*_ ∼12.6 μM), demonstrating target-specific binding. **B)** Concentration-dependent binding of anti-CD25 aptamers to CD25-positive Ramos cells (green) and CD25-negative HEK293T cells (blue). Aptamer 77 (left) exhibits high-affinity binding (*K*_*d*_ ∼31 nM), whereas Aptamer 64 (right) shows moderate affinity (*K*_*d*_ ∼2.7 μM), with minimal binding to CD25-negative cells. **C)** Representative flow cytometry histograms of purified CD4^+^ T cells treated with daunorubicin-loaded Aptamer 77, Aptamer 64, a CD117-targeting control aptamer, or left untreated showing selective response in CD25+ cells. **D)** Background-subtracted MFI quantification corresponding to panel **C**, showing selective depletion of CD25^+^ cells following treatment with Aptamer 77 (***P < 0.001; ns, not significant). **E)** Representative flow cytometry histograms of Raw264.7 cells treated with daunorubicin-loaded aptamers or control, demonstrating no selective response in CD25-negative cells. **F)** Background-subtracted MFI quantification for Raw264.7 cells corresponding to panel **E**, showing no significant differences between treatments (ns, not significant). **G)** Representative flow cytometry histograms of PBMCs treated with daunorubicin-loaded aptamers, shown for CD25^−^, CD25^+^, and CD4^+^CD25^+^ subpopulations. **H)** Background-subtracted MFI quantification corresponding to panel **G**, demonstrating selective depletion in CD25^+^ and CD4^+^CD25^+^ populations following treatment with Aptamer 77 (*P < 0.05; ***P < 0.001).

Similarly, anti-CD25 Aptamers 64 and 77 were evaluated under identical conditions in CD25-positive Ramos cells and CD25-negative HEK293T cells (**Figure 2B, S8**). Both aptamers demonstrated robust, concentration-dependent binding to Ramos cells, with negligible binding to HEK293T cells. Binding analysis revealed K_d_ values of approximately 2.7 μM for Aptamer 64 and 31 nM for Aptamer 77, identifying Aptamer 77 as a high-affinity CD25 binder.

To evaluate functional activity beyond target binding, we conjugated Aptamers 64 and 77 with the cytotoxic DNA-intercalating agent daunorubicin. Single daunorubicin molecules intercalate within duplex regions of the aptamer without compromising target recognition. Treatment of purified CD4^+^ T cells with daunorubicin-loaded Aptamer 77 resulted in selective depletion of the CD4^+^CD25^+^ subpopulation (**Figure 2C,D; Figure S9**), while CD4^+^CD25^−^ cells exhibited approximately 2.5-fold lower depletion, demonstrating selective payload delivery. In contrast, Aptamer 64 did not elicit comparable cytotoxicity, likely due to the assay concentration (1000 nM), which exceeded the K_d_ of Aptamer 77 by >30-fold but remained below the optimal binding range for Aptamer 64. A control daunorubicin-loaded aptamer targeting CD117, a marker not expressed on CD4^+^ T cells, produced minimal and comparable cytotoxicity across CD25^+^ and CD25^−^ populations. Consistently, CD25-negative Raw264.7 cells showed no loss in viability upon treatment (**Figure 2E,F; Figure S10**). To further stringently assess specificity, PBMCs differentiated under T-cell polarizing conditions were treated with daunorubicin-loaded aptamers (**Figure 2G,H; Figure S11**). Aptamer 77 again selectively depleted CD4^+^CD25^+^ cells while exerting minimal effects (<2-fold) on CD4^+^CD25^−^ or double-negative populations. The CD117-targeting control aptamer showed negligible cytotoxicity across all subpopulations.

AptaBLE represents a significant advance in computational aptamer development. By operating directly on sequence data, our model circumvents the scarcity of structural information while capturing the complex interaction patterns critical for aptamer-protein binding. Our framework’s capability to generate novel aptamer sequences with desired binding properties opens new possibilities for rational aptamer design, potentially reducing the time and resources required for aptamer development. AptaBLE has the potential to significantly accelerate the development of aptamer-based therapeutics and diagnostics by rational streamlining the discovery process.

## Author Contributions

S.P., K.F, and S.Y conceived the idea and guided the project. S.P. performed the development and training of the AptaBLE model with input from F.Z.P, P.C., S.Y, and K.F. K. F. performed structural characterization of the generated aptamers. D.G. and A.D. performed all in vitro experiments, including flow cytometry and functional assays with supervision from X.Y. D.G and A.D performed analysis of data generated from in vitro experiments. S.P. led writing of the manuscript with contributions from all authors. All authors have read the paper, agreed to its content and approved the final submission.

## Declarations

The authors acknowledge that this work was funded by Atom Bioworks, Inc.

## Acknowledgements

The authors would like to thank Shrey Shah and Meenakshi Ranasinghe for consulting on the project.

## Materials and Methods

### Architecture

AptaBLE employs a symmetric architecture that processes both protein and aptamer sequences through specialized pretrained encoders, followed by a fusion mechanism for combining embeddings into a shared aptamer-protein representation (**Figure 1A**). It consists of symmetric cross-attention modules that enable bidirectional information flow between protein and aptamer representations, capturing complex interaction patterns without providing any structural information. The fusion architecture comprises two cross-attention modules and two self-attention modules, the outputs of which are summed, pooled, and concatenated to arrive at a joint representation. This is ultimately processed by a multi-layer perceptron to generate binding predictions.

#### Protein encoder

We use ESM-2^11^ with 150M parameters, pretrained on 250 million protein sequences from UniRef50. The encoder processes protein sequences through 33 transformer layers with 1280-dimensional hidden states, generating per-residue embeddings. We freeze all encoder parameters during training to preserve evolutionary information while reducing computational cost and overfitting risk.

#### Aptamer encoder – DNA

For DNA aptamers, we use Nucleotide Transformer v2-50M^12^ (nucleotide-transformer-v2-50m-multi-species), pretrained on 850 billion nucleotides from reference genomes of 3200 species. The encoder uses 6-nucleotide tokenization (6-mers), processing sequences through 12 transformer layers with 512-dimensional hidden states. All parameters are frozen during training.

#### Aptamer encoder – RNA

For RNA aptamers, we use RNA-FM^13^, pretrained on 23 million RNA sequences from RNAcentral. The encoder processes single nucleotides through 12 transformer layers with 640-dimensional hidden states. Parameters are frozen during training.

#### Dimension alignment

To enable cross-attention between protein (1280-dim) and aptamer (512-dim for DNA, 640-dim for RNA) embeddings, we apply learned linear projections to align dimensions. For DNA models, we project protein embeddings from 1280 to 512 dimensions. For RNA models, we project protein embeddings to 640 dimensions. These projections use standard linear layers without bias: 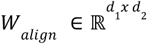 where *d*_1_ = 512, *d*_2_ = 1280 for DNA and *d*_1_ = 640, *d*_2_ = 1280 for RNA.

#### Fusion Mechanism

As shown in analogous work^25^, we leverage a fusion mechanism to combine embeddings of the aptamer and protein modalities. We take a vectorized protein *P* ∈ *W*^*m*^ of length *m* and aptamer *A* ∈ *W*^*n*^ of length n and map the pair to a score between 0 and 1.

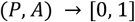

To do this, we leverage two pretrained encoders to produce embeddings of the protein 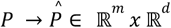 and the aptamer 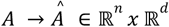. These are subsequently fused via a cross attention fusion module to generate a combined representation of the pair. For each cross attention fusion module, we learn three weight matrices: *W*_*q*_, *W*_*k*_, *W*_*v*_ ∈ *R*^*hxh*^ which are used to generate the queries, keys and values, or *Q, K, V*, for the module. We therefore generate the following matrices:

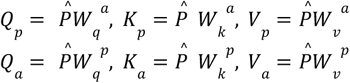

We also learn another set of weight matrices for the self-attention module:

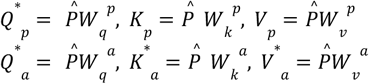

From these matrices, we generate the fused representations as below:

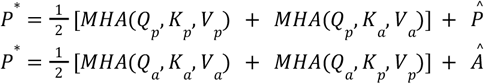

To combine *P*^*^ ∈ *R*^*m,d*^ and *A* ∈ *R*^* *n,d*^, we mean-pool each matrix and concatenate the result to yield a vector *V*_0_ ∈ *R*^2*d*^. This vector is finally downsized by a series of linear-ReLU-batchnorm modules until a prediction score is produced via a sigmoid activation.

### De Novo Generation Algorithms

To generate aptamers *de novo*, we adopt and extend two methods inspired by multi-objective conditional sequence design where AptaBLE is used to compute rewards, and navigate sequence-space exploration with either randomly-generated sample sequences or diffusion-generated sample sequences.

#### Monte Carlo Tree Search

Our first approach is a simple Monte Carlo Tree Search method^23^ only requiring the use of the AptaBLE classifier. In addition to the below, the full algorithm is described in Algorithm 1.

##### Initialization

We take our final aptamer as a sequence of fixed length *N*, where initially the sequence is completely empty. At each iteration of the MCTS, a new token is either appended or prepended to the existing aptamer sequence based on the selection-expansion-rollout cycle followed by the algorithm.

##### Selection

From the root node, the algorithm progresses down the MCTS tree based on the UCT (upper confidence bounds applied to trees) score, defined as the following:

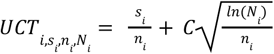

where *i* identifies the node, *s*_*i*_ is the accumulated exploitation reward for node *i* calculated during backpropagation, *n*_*i*_ is the number of visits to node *i, N*_*i*_ is the number of visits to the parent node of node *i*, and *C* is a tunable exploration parameter set to 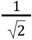 according to prior work^35^. Ultimately, a leaf node is reached following the optimal UCT path.

##### Expansion

From the selected leaf node, a new child node is added. This reflects either the appending or prepending of a nucleotide base to the existing sequence represented by the currently selected node (i.e. _A, _C, _G, _T, A_, C_, G_, or T_, where _ denotes the current state of the node).

##### Rollout

For each child node 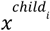, the remainder of the aptamer sequence is completed by randomly appending nucleotides from the model alphabet until the sequence reaches length *N*. This random sequence generation from the child node denotes one sample from the child node state. Ultimately, each sequence generated from this method reflects the child node’s construction and is passed into AptaBLE alongside the target protein for score computation. This reflects an individual play-out.

##### Backpropagation

For each child node 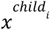, we update its cumulative reward denoted by its UCT by populating the *s*_*i*_value using the AptaBLE scores corresponding to that play-out. This procedure of exploring random play-outs is repeated for the indicated number of iterations in the MCTS. Ultimately, the procedure repeats from the selection step, with the child node having the greatest UCT score being added to the ensuing selection path.

##### Reward

The reward vector for a node consists of two scores which equally contribute to the *s* _*i*_ value in the UCT computation. The first is the AptaBLE score against the target protein and the second is the AptaBLE score against an off-target protein subtracted from 1 to function as a ‘negative’ selection. These scores are equally weighted and combined, then serving as the sequence’s exploitation reward within its UCT.

#### Monte Carlo Tree Guidance

Our second approach is applying Monte Carlo Tree Guidance^24^ to aptamer sequence design. This method requires the use of the AptaBLE classifier and an appropriate masked diffusion model to sample sequences from each explored state. In addition to the below, the full algorithm is described in Algorithm 2.

Here, *de novo* generation requires the use of a masked diffusion model (MDM) for sampling sequences at the rollout stage. We use a pretrained RNA masked diffusion model^26^ for sampling sequences from each ssDNA aptamer state.

##### Initialization

We initialize a fully masked tokenized sequence with length *L*. This becomes the root node of the MCTS tree with timestep *t* = 0. Additionally, we construct a vector scoring function *f*: *V*^*L*^ → *R*^*K*^ that returns a score vector given a clean input sequence *x*_0_ ∈ *V*^*L*^ and initialize an empty Pareto-optimal set *P*^*^ = {}. We define the following hyperparameters: number of MCTS iterations *N*_*iter*_ and number of children nodes *N*_*child*_. The root node is initialized as a fully-masked sequence of length *L*.

##### Selection

To determine the next unmasking step from a parent node with an already expanded set of child sequences 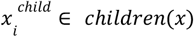, we compute a selection score vector 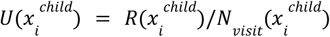 that divides the cumulative reward vector 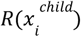 of a child node to the number of times it was visited 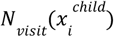. This encourages exploration of child nodes that have not been visited, in addition to high-reward nodes. Then, we select from the Pareto-optimal set of child nodes based on the normalized selection score vector.

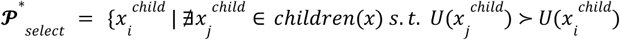

If we select a non-leaf node, we restart the selection process with the selected node as the new parent until we reach an expandable leaf node or a fully unmasked sequence. In the case of a fully unmasked sequence, we restart the selection from the root node.

##### Expansion

From the leaf node *x*_*t*_ at time *t*, we sample *N*_*child*_ distinct sequences by applying Gumbel noise to the denoising distribution 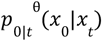

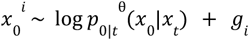

where *g*_*i*_ ∼*Gumbel*(0, 1). Then, for each child node 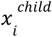, we randomly select *k* positions in *x*_*t*_ to unmask according to 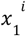.

##### Rollout

For each child node 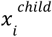, we use self-planned ancestral sampling to obtain a fully unmasked sequence 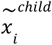. This is in stark contrast to the aforementioned MCTS method, where sequences corresponding to each state are produced completely at random. Then, we feed each of the clean sequences into the vector score function 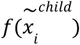 to obtain a score vector. We then compute the reward vector 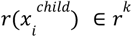 of each child 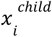 where each element is the fraction of sequences in the Pareto-optimal set *x*^*^∈ *P*^*^ that 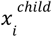 dominates.

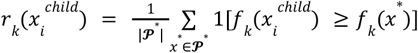

For all *N*_*child*_ children sequences, we add 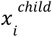 to *P*^*^ if it is *non-dominated* by all sequences *x*^*^ ∈ *P*^*^ and remove all sequences in *P*^*^ dominated by some 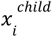.

##### Backpropagation

For each child node 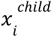, we use its reward vector 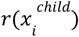 to initialize the cumulative reward 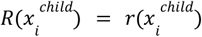 and the number of visits 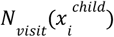 is set to 1.

We then trace backward from 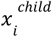 through its ancestors up to the root node 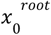, updating each node *x* along the path by accumulating the child rewards and incrementing the visit count:

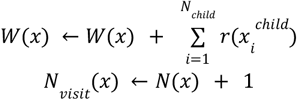

These cumulative updates influence future selection, prioritizing unmasking paths that lead to higher-reward sequences for further expansion. After *N*_*iter*_ iterations of these four steps, we return the set of aptamer sequences optimized across the set of properties *x*^*^ ∈ *P*^*^ and their corresponding score vectors *f*(*x*)^*^.

##### Reward Vector

Similar to the MCTS method, the reward vector here reflects the AptaBLE scores for each aptamer against the target protein and an off-target protein serving as an *in silico* negative control (subtracted from 1). These scores are equally weighted and combined, forming the backpropagated reward.

#### Settings

All generated libraries consisted of 100 40N ssDNA aptamers.

##### MCTS

- Number of iterations: 100
- Rollout strategy: Random nucleotide completion from each leaf node
- Number of children: 8

##### MCTG

- Number of iterations: 128
- Rollout strategy: masked diffusion model sampling
- Number of children: 12

##### Training Dataset

###### Positive examples

We compiled aptamer-protein binding pairs from three sources:

1. UTexas Aptamer Database^27^: 847 DNA aptamers, 423 RNA aptamers with validated binding
2. Li et al. 2014 dataset^28^: 256 additional RNA aptamers
3. Proprietary data from Atom Bioworks

After removing duplicates and filtering for sequence quality (no ambiguous bases, length 20-100nt), the final positive dataset contained: 1,089 DNA aptamers binding 127 protein targets, 757 RNA aptamers binding 94 protein targets. We note here that neither hCD117 nor any hCD117 aptamers were included in our model training dataset.

###### Negative examples

For each positive aptamer, we generated four negative examples by shuffling the sequence while preserving nucleotide composition and length. Shuffling was performed using Python’s random.shuffle with different random seeds. We verified that shuffled sequences have different predicted secondary structures (>50% difference in base pairing) using ViennaRNA RNAfold^29^.

This 4:1 negative:positive ratio was chosen based on preliminary experiments showing that higher ratios improved validation performance without requiring prohibitive computational resources. The shuffling strategy creates chemically plausible sequences (matching length and composition of real aptamers) while eliminating binding, forcing the model to learn sequence-structure-function relationships rather than simple nucleotide frequencies.

###### Dataset splitting

To prevent data leakage and test generalization to novel proteins, we performed homology-based splitting:

- Cluster all protein targets using MMseqs2^30^ with parameters: --min-seq-id 0.8, --cov-mode 0, -c 0.8
- Assign entire clusters to training (80%), validation (10%), or test (10%) sets
- This ensures no test protein shares >80% sequence identity with any training protein

#### Benchmark Dataset

We collected an additional benchmark set of 16 ssDNA aptamers and 39 RNA aptamers to use for evaluating models capable of predicting aptamer-protein interactions. These aptamers and their protein targets are omitted from the training dataset. We evaluate three models, including AptaBLE, on each of these datasets in Figure S1A,B.

#### Auxiliary Benchmark Set

This dataset consists of four known-binding ssDNA aptamer-protein pairs which we highlight in this work – two aptamers binding *SARS-CoV-2* Spike glycoprotein^31^, one aptamer binding gp120^32^, and the final aptamer binding basic fibroblast growth factor^33^. Results on these pairs are reported in Figure S1C.

##### Retrospective Analysis

Aptamers from the described hCD117 SELEX were ultimately clustered by FastAptamer^9^ and retrospectively scored via AptaBLE to identify which clusters are better predicted to serve as functional binders to hCD117 (**Figure S2C**). It is expected that the higher-scoring clusters would have been prioritized in the analysis rather than simply advancing sequences based on their number of copies following sequencing. As sequences from clusters 8-11 were not characterized for binding, focus was kept on clusters 1-7. From **Figure S2A**, sequences with the highest copy number are not always scored the highest by AptaBLE within their own cluster. Additionally, there is a tight correspondence between binding affinity from a representative aptamer within each cluster and the overall distribution of AptaBLE scores within a cluster (**Figure S2C**). From this analysis, sequences from clusters 2, 3, and 7 would have been prioritized for characterization using a workflow involving AptaBLE. Notably, the constituent aptamers from these clusters had the strongest binding affinity to hCD117 as determined via biolayer interferometry.

Secondary structure prediction for all aptamers was done with ViennaRNA^29^.

##### Biolayer Interferometry

Binding affinity measurements for all synthesized aptamers shown in **Figure 2A** were performed in a biolayer interferometry (BLI) assay with an Octet GatorPrime instrument. All assays were performed at 30°C in 96-well BLI 96-Flat Black Plates, with samples prepared in an assay buffer containing 50 mM HEPES, 150 mM NaCl, and 5 mM MgCl_2_ (pH 7.4). For kinetic analysis of all free hCD117 aptamers (ABWUC1-4, ABW02-07), anti-VHH biosensors were used to immobilize hCD117 (Sino Biological, Inc.) at 50 µg/mL for 180 seconds, followed by capture of 6x-His-tagged recombinant human CD4 (rCD4; Sino Biological, Inc.) at 3 µg/mL for 180 seconds. All aptamers were then tested at concentrations of 20,000, 10,000, 5,000, 2,500, 1,250, and 625 nM, with association and dissociation phases of 900 seconds each.

##### Cell Maintenance

Ramos, RAW 264.7, and HEK293T cell lines were procured from the American Type Culture Collection (ATCC; CRL-1596, TIB-71, CRL-3216). All cell lines were cultured in RPMI-1640 medium supplemented with 10% fetal bovine serum (FBS) and 1× antibiotic–antimycotic solution. Primary cells, including CD4^+^ T cells and peripheral blood mononuclear cells (PBMCs), were obtained from ATCC (PCS-800-016, PCS-800-011) and cultured in ImmunoCult medium supplemented with 10% FBS, 1× antibiotic–antimycotic solution, interleukin-2 (IL-2), and human CD3/CD28/CD2 T-cell activators. All cells were maintained in a humidified CO_2_ incubator at 37 °C with 5% CO_2_ and were passaged upon reaching approximately 70% confluence. For experimental use, cells were harvested by treatment with trypsin-EDTA, followed by centrifugation at 500 × g for 10 minutes at room temperature. The resulting cell pellet was resuspended in the appropriate culture medium. For each experiment, a minimum of 10,000 cells per replicate was used.

##### Flow Cytometry

To assess the specificity and affinity of FAM-labeled aptamers (Aptamer 77 selected for CD25 using AptaBLE) toward Ramos cells (CD25^+^), a comparative binding assay was performed using HEK293T cells (CD25^-^) as a negative control. Aptamer-FAM was first dissolved in nuclease-free water to a final concentration of 1 mM. A 50 µM working solution was prepared using a folding buffer containing 20 mM HEPES (pH 7.4), 150 mM NaCl, and 5 mM MgCl_2_. Aptamers were folded by heating at 95 °C for 5 min, cooling at 4 °C for 5 min, incubating at 25 °C for 5 min, and holding again at 4 °C for 5 min. The folded aptamers were serially diluted 4-fold in 1× selection buffer, ranging from 50 µM to 0.781 µM.

Ramos cells were harvested from a T-25 flask and transferred to a 15 mL centrifuge tube. The cells were pelleted by centrifugation at 400 × g for 10 min, after which the supernatant was completely removed. The pellet was gently resuspended in 1 mL of 1× selection buffer (20 mM HEPES (pH 7.4), 150 mM NaCl, 5 mM MgCl_2_ and 10% FBS). Cell concentration and viability were determined using Trypan Blue. HEK293T cells were cultured and prepared in parallel, to serve as the negative control group.

A 96-well plate was prepared for the binding assay. Each well contained a total of 10,000 cells in a final volume of 100 µL. To achieve this, an appropriate volume of 1× selection buffer was first added to each well so that, after cell addition, the total volume equaled 90 µL. The calculated volume of cell suspension corresponding to 10,000 cells was added to each well. 10 µL of each concentration of folded aptamer was added to the designated wells with six replicates per concentration. Wells containing either Ramos or HEK293T cells without aptamer were used as fluorescence background controls. The plate was gently mixed and incubated at 37 °C in a CO_2_ incubator for 40 min.

After incubation, the plate was centrifuged at 450 × g for 10 min at room temperature. The supernatant was removed using a multichannel pipette. Each well was resuspended in 200 µL of 1× selection buffer.

Flow cytometry was performed using an Attune cytometer equipped with a 488 nm laser and BL1 channel to detect FAM fluorescence. Standard gating was applied to select single, viable cells based on FSC-A vs SSC-A and FSC-H vs FSC-A parameters. A minimum of 5,000–10,000 events were collected per sample. Binding efficiency was quantified by comparing the median fluorescence intensity (MFI) in Ramos versus HEK293T populations. All experiments were performed in six replicates, and K_d_ was determined by fitting the MFI values against aptamer concentration using a one-site specific binding model. Nonlinear regression analysis was carried out using GraphPad Prism to obtain best-fit parameters and standard errors.

##### Aptamer Cell Binding Assays

For cell binding studies, fluorophore-tagged aptamers (obtained from IDT, USA) were tested at multiple concentrations, with appropriate dilutions prepared independently for each concentration. A minimum of six biological replicates were used per cell type, with each replicate containing at least 10,000 cells. Prior to incubation, aptamers were folded as described earlier (heating to 95 C and rapid cooling on ice) and then added to the cells in the appropriate binding buffer. Cells were incubated with aptamers for 1 h in a humidified CO_2_ incubator. Following incubation, cells were collected by centrifugation at 500 × g for 10 min and resuspended in the same binding buffer. Cells were subsequently washed twice under identical conditions to remove unbound aptamers. After the final wash, samples were processed for flow cytometry analysis. Mean fluorescence intensity (MFI) obtained from fluorophore-tagged aptamer was used to assess binding efficiency for each aptamer and cell type. MFI values across aptamer concentrations were further analyzed to characterize binding dynamics and were used to calculate the equilibrium dissociation constant (KD) for each aptamer.

##### Aptamer Functional Assays

For functional studies, aptamers were first loaded with the cytotoxic drug daunorubicin using previously described methods. Daunorubicin intercalates into double-stranded DNA, inducing DNA strand breaks and resulting in cell death. Briefly, aptamers were folded as described above and incubated with daunorubicin overnight to allow drug loading. Unbound daunorubicin was subsequently removed by centrifugation using a membrane-based centrifugal filtration unit. Daunorubicin-loaded aptamers were then incubated with cells for 18 h under standard culture conditions. Following treatment, cells were collected and subjected to flow cytometry–based analysis. For immunophenotyping, cells were stained with fluorophore-conjugated antibodies targeting CD4 and/or CD25 for 30 min using an appropriate flow cytometry staining buffer. For apoptosis analysis, cells were incubated with fluorophore-conjugated Annexin V in Annexin V–specific binding buffer containing calcium for an additional 30 min. After each staining step, cells were washed at least three times to remove unbound antibodies or Annexin V conjugates. Cells were subsequently analyzed by flow cytometry to assess cell depletion and apoptotic responses.

For CD4^+^ T cells and RAW cells, samples were stained only with a fluorophore-conjugated CD25 antibody. Cell populations were gated into CD25^+^ and CD25^−^ subsets, and Annexin V signals were quantified for the total (bulk) cell population as well as separately for the CD25^+^ and CD25^−^ populations. For PBMC samples, cells were stained with two fluorophore-conjugated antibodies targeting CD4 and CD25. Annexin V staining was subsequently analyzed for distinct subpopulations, including CD4^+^CD25^+^ double-positive cells, CD4^+^CD25^−^ cells, and the bulk CD25^+^ population, in addition to the total PBMC population. This gating strategy enabled subpopulation-specific assessment of apoptosis following aptamer-mediated cell depletion. MFI values across various populations for Annexin signals were further utilized to quantify relative cell death across cell and aptamer types.

## Appendix

### AptaBLE performance exceeds competing methods

AptaBLE was trained on validated, published aptamers^27,28^, validated proprietary aptamers, and artificially generated pseudo-negative derivatives of those aptamers. Further details regarding our training and benchmarking dataset and implementation can be found in the benchmark and auxiliary benchmark sections.

We benchmarked AptaBLE against two representative approaches: AptaTrans^14^, the current state-of-the-art sequence-based method, and AlphaFold3^20^, leveraging structure-based prediction. AptaBLE significantly outperformed both baselines across all metrics for DNA aptamers (**Figure S1A**). The comparably high recall is particularly important for SELEX analysis, as false negatives represent missed therapeutic candidates. For RNA aptamers, AptaBLE also performed favorably over the competing models, though the margin over AptaTrans was smaller (**Figure S1B**). AptaBLE additionally outperformed both models in predicting stable ssDNA aptamer-protein complexes in a small auxiliary dataset (**Figure S1C**).

Runtime analysis demonstrated AptaBLE’s practical efficiency. Scoring a single aptamer-protein pair required 48 milliseconds on average in batch mode on an A100 GPU,

### Fusion attention maps suggest potential aptamer-protein binding interfaces

To investigate the model’s representation of both nucleic acids and amino acids, we studied attention maps generated during inference on two aptamer-protein complexes with known crystal structures. These aptamers, RBD-1C and RBD-4C, bind to the *SARS-CoV-2* Spike protein receptor binding domain (RBD)^31^.

We observe one of our attention heads in the cross attention fusion module representing protein-attended nucleic acids reflecting atomic interactions seen at the binding interface between the S-protein and the aptamer (**Figure S2A,B**). Notably, we find that each nucleic acid implicated in the binding interface is ‘more’ attended to in both cases. However, we do not always see that the corresponding amino acid participating in the hydrogen bonding interaction also is relatively attended to within that column. As such, it is inconclusive as to whether these cross attention maps indicate inter-residue interaction.

### *De Novo* Generation

For cell surface receptor targets human CD25 and TIGIT, two independent aptamer libraries were generated using AptaBLE-MCTS and AptaBLE-MCTG. Both libraries spanned 100 40N aptamers in total and were designed with a prompt for optimizing AptaBLE score for the intended target protein (either CD25 or TIGIT) and optimizing (1 - AptaBLE score) for human albumin. The latter was provided for negative selection. Aptamers were selected for synthesis by identifying structurally distinct aptamers following a hierarchical clustering based on predicted secondary structure (via ViennaRNA) and following structure prediction via Chai-1 Version 0.6.1 with refined docking performed with HADDOCK 3.0 (described below in **Docking**).

Aptamers with the highest ipTM and LIS score were selected for synthesis after verifying that their predicted structure dot-bracket edit distances differed by more than 10. 3 aptamers were synthesized from each library. In total, 5 aptamers were synthesized each for CD25 and TIGIT (3 derived from MCTS-generated library, 2 from MCTG-generated library). We include results here for only the top-2 binding aptamers to each protein target following a cell-based flow cytometry assay (described in Methods: **Flow Cytometry**). Aptamers 77 and and 27 were produced via MCTS while 64 and 22 were produced via MCTG.

### Docking

To evaluate binding of both wild type and designed aptamers to these targets *in silico*, We used HADDOCK3 (version 2024.10.0b7; Giulini Et Al., 2025) for modeling biomolecular complexes. PDB structures were initially predicted using Chai-1. Docking simulations were performed using a workflow consisting of topology generation (topoaa module), rigid-body docking (rigidbody module), flexible refinement with semi-flexible simulated annealing (flexref module), energy minimization (emref module), and fraction of common contacts (FCC) clustering (clustfcc module with default clustcutoff = 0.6). Models within each cluster were ranked according to the HADDOCK score, a weighted energy function combining van der Waals energy, electrostatic energy, desolvation energy, buried surface area, and ambiguous interaction restraint terms. For each aptamer-protein complex, the cluster exhibiting the median HADDOCK score was identified, and the top-ranked model from this cluster was selected as a snapshot of the complex interaction. Results were visualized with ChimeraX^35^.

#### Environment

We train and perform our experiments on a single a2-ultra-gpu machine type available via the Google Cloud Platform. This machine has 170 GB RAM across 12 vCPUs, an Intel Cascade Lake processor, and an NVIDIA A100 80gb GPU. We implement our model using Python 3.9 and PyTorch 2.3.0.

However, we note that our model could be trained on any compute platform supporting a GPU with at least 48gb memory.

#### Training

##### Loss function

Binary cross-entropy loss:

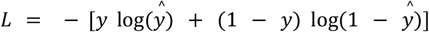

where y ∈ {0,1} is the true label and ŷ ∈ [0,1] is the predicted probability.

##### Optimization

AdamW optimizer with parameters: β_1_=0.9, β_2_=0.999, ε=10^−8^, weight decay=0.01. Initial learning rate: 10^−5^. Learning rate schedule: MultiStepLR with decay factor 0.1 at epochs [5, 10, 20].

##### Training hyperparameters

- Batch size: 16 (limited by GPU memory for long sequences)
- Maximum epochs: 30 (DNA), 20 (RNA)
- Gradient clipping: max norm 1.0
- Mixed precision training: FP16 using torch.cuda.amp
- Early stopping: patience of 5 epochs on validation AUC-ROC

##### Hardware and runtime

All training performed on Google Cloud Platform a2-ultragpu-1g instances with NVIDIA A100 80GB GPU, 12 vCPUs, 170GB RAM. Typical training time: 18 hours for DNA model, 14 hours for RNA model. Memory usage: 65GB GPU memory at peak (during backpropagation for longest sequences).

We use the MultiStepLR Scheduler with an initial learning rate of 1 * 10^−5^ which decays by 0.1 every 5 timesteps. We use AdamW as our optimizer and binary cross-entropy as our loss function, implemented by PyTorch. Our batch size is fixed to 16 across both versions of our model. We performed a hyperparameter sweep over the initial learning rate, batch size and random seed for model configuration. We test several checkpoints across several trained models, of which we report the best-performing results here as determined purely through benchmark dataset performance.

##### Baseline comparisons

###### AptaTrans

We implemented AptaTrans^14^. We retrained the model using their published code and dataset for fair comparison. For DNA aptamer evaluation, we converted sequences to RNA (T→U substitution) as required by their RNA-specific encoder. We used their best checkpoint (epoch 50) for all evaluations.

###### AlphaFold3

We ran AlphaFold3^21^ structure prediction using the default server (version 2024.1) on a subset of 200 test examples due to computational cost (5 min/prediction). We used predicted local distance difference test (pLDDT) scores >70 as indicating binding. For aptamer-protein pairs with multiple predicted structures (5 models per prediction), we used the highest-confidence model.

###### Statistical testing

All performance comparisons used bootstrap resampling (1000 iterations) to compute 95% confidence intervals. Significance was assessed using permutation tests (1000 permutations) with α=0.05.

###### Structure data

We obtained crystal structures for RBD-1C (PDB: 7KMD) and RBD-4C (PDB: 7KME) from Lin et al. 2022^31^. Structures were processed using BioPython to extract coordinates.

###### Contact definition

Protein-aptamer contacts were defined as residue-nucleotide pairs with any heavy atom within 5Å distance. This yielded 47 contacts for RBD-1C and 52 contacts for RBD-4C. We also note that each column in the attention map reflects a disjoint group of 6 nucleic acids, as we follow the aptamer encoder’s 6-mer tokenization scheme.

###### Attention extraction

For each aptamer-protein pair, we:

1. Performed forward pass through AptaBLE
2. Extracted attention weights from the protein-to-aptamer cross-attention module (8 heads, each L_p_×L_a_ matrix)
3. Averaged across attention heads to yield single attention matrix A ∈ ℝ^(L_p_×L_a_)
4. For Nucleotide Transformer’s 6-mer tokens, aggregated attention to nucleotide level by assigning each nucleotide the maximum attention from any overlapping token

##### Architecture

We list our model architecture configurations here.

**Cross-attention module**: The hidden size and number of heads are 512 and 8..

**Self-attention module**: The hidden size and number of heads are 512 and 8.

**Reshape layer**: We reshape ESM2 embeddings to have a hidden dim of either 512 or 640 from 1280 for compatibility with Nucleotide-Transformer or RNA-FM, respectively.

**MLP**: We pass the fused representation through an MLP with linear layers mapping from either 1024 or 1280 (depending on DNA or RNA, respectively), to 512, to 256, to

1. Each layer is followed by a ReLU activation and 1-D Batchnorm.

##### Algorithms

Here, we provide algorithms for both aptamer generation methods described in this work.

###### Algorithm 1

Monte-Carlo Tree Search generation of single-stranded DNA aptamers.

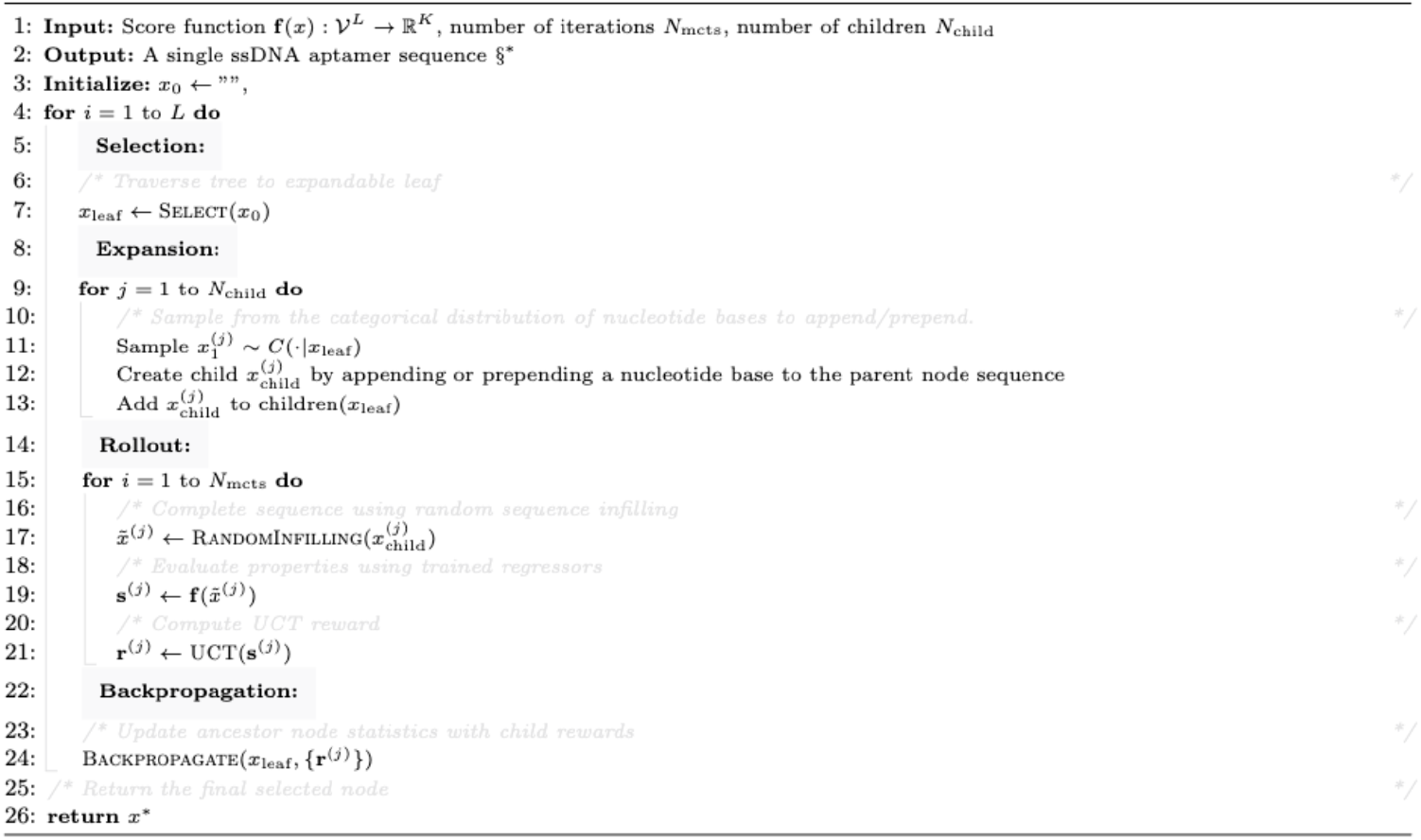

**Algorithm S1. Monte Carlo Tree Search.** Algorithm adapted from [23]

###### Algorithm 2

Monte-Carlo Tree Guidance generation of single-stranded DNA aptamers.

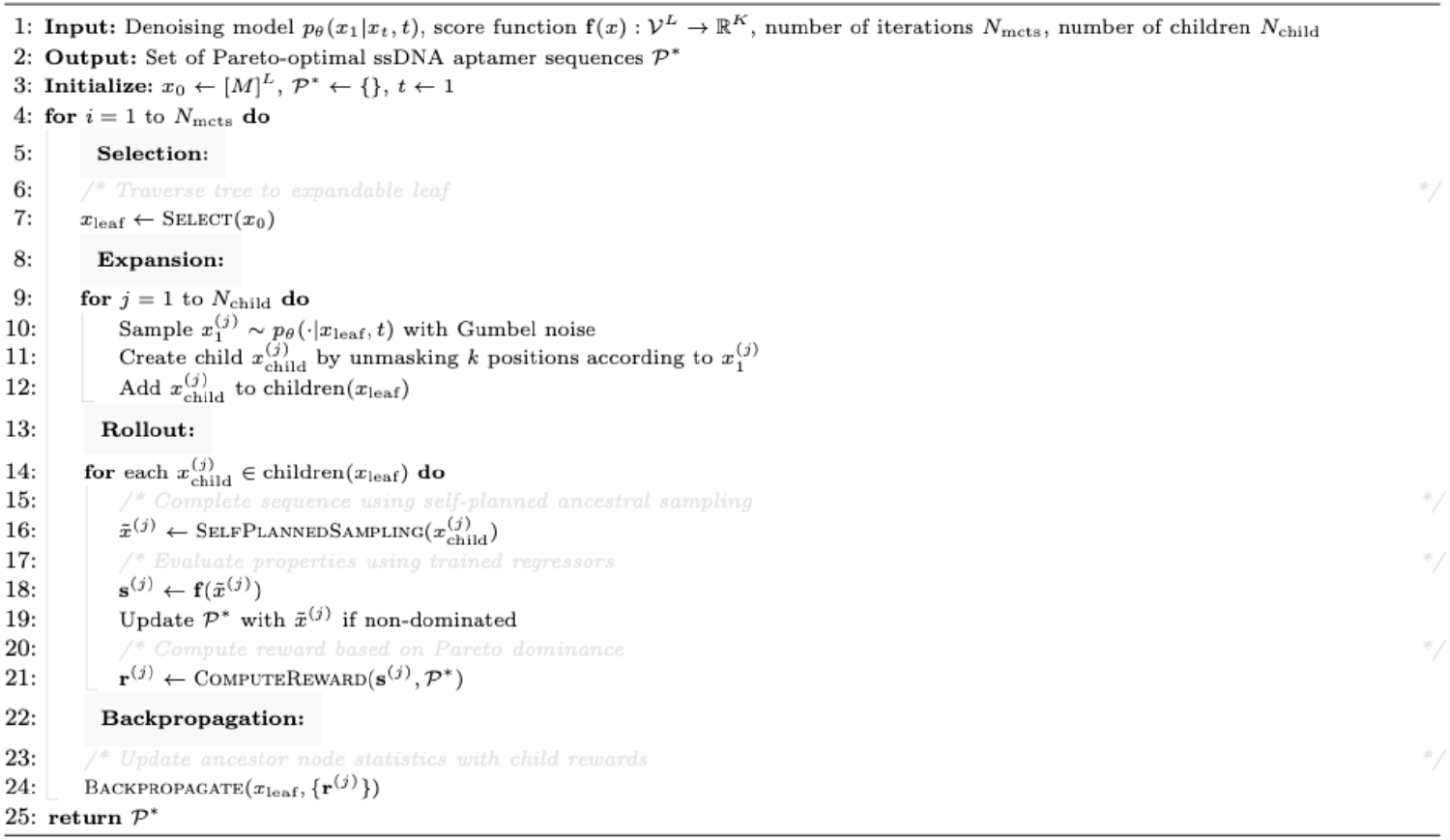

**Algorithm S2. Monte Carlo Tree Guidance.** Algorithm adapted from [24].

---

**Table S1:**
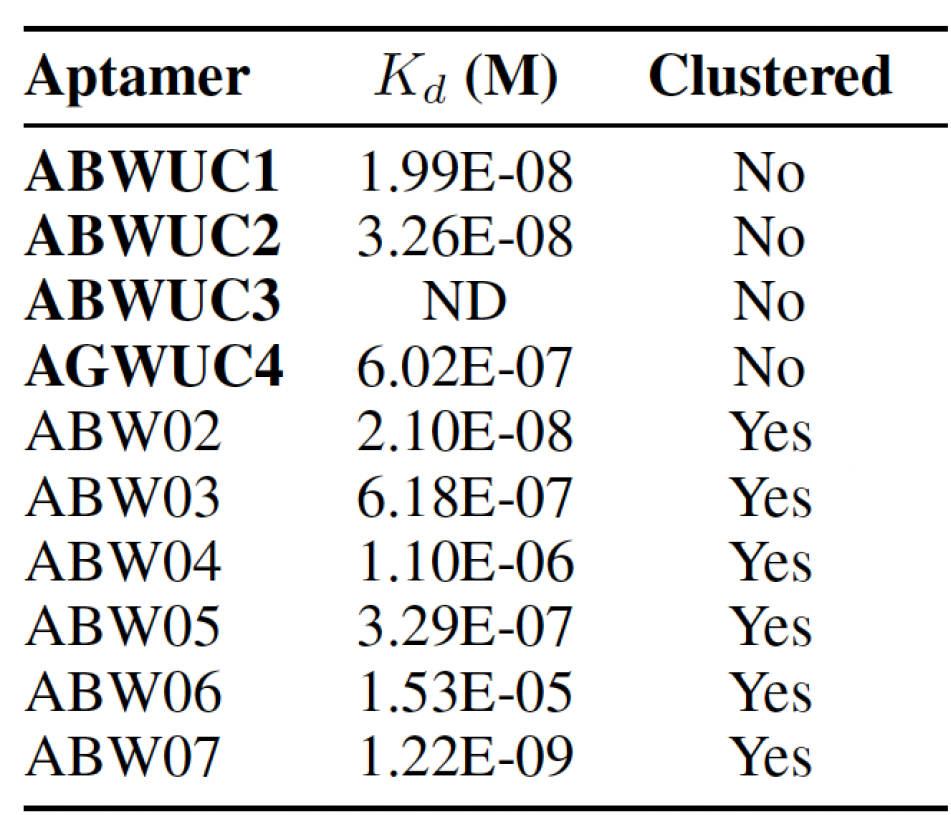
Aptamers discovered to bind human CD117 following 9 Rounds of SELEX. Aptamer library was previously filtered and clustered via FastAptamer to identify highly-enriched sub-populations of ssDNA sequences. Clusters were assigned leading aptamers having a high copy number following sequencing and all aptamers belonging to a cluster possessed a sequential edit distance less than or equal to 3 to the lead. Aptamers that were unclustered were dismissed from further consideration during initial discovery phase. Retrospective analysis using AptaBLE discovered several aptamers binding to CD117 (**bold**).

| **Aptamer** | $K_d$ (M) | **Clustered** |
| --- | --- | --- |
| **ABWUC1** | 1.99E-08 | No |
| **ABWUC2** | 3.26E-08 | No |
| **ABWUC3** | ND | No |
| **AGWUC4** | 6.02E-07 | No |
| ABW02 | 2.10E-08 | Yes |
| ABW03 | 6.18E-07 | Yes |
| ABW04 | 1.10E-06 | Yes |
| ABW05 | 3.29E-07 | Yes |
| ABW06 | 1.53E-05 | Yes |
| ABW07 | 1.22E-09 | Yes |

**Figure S1.**
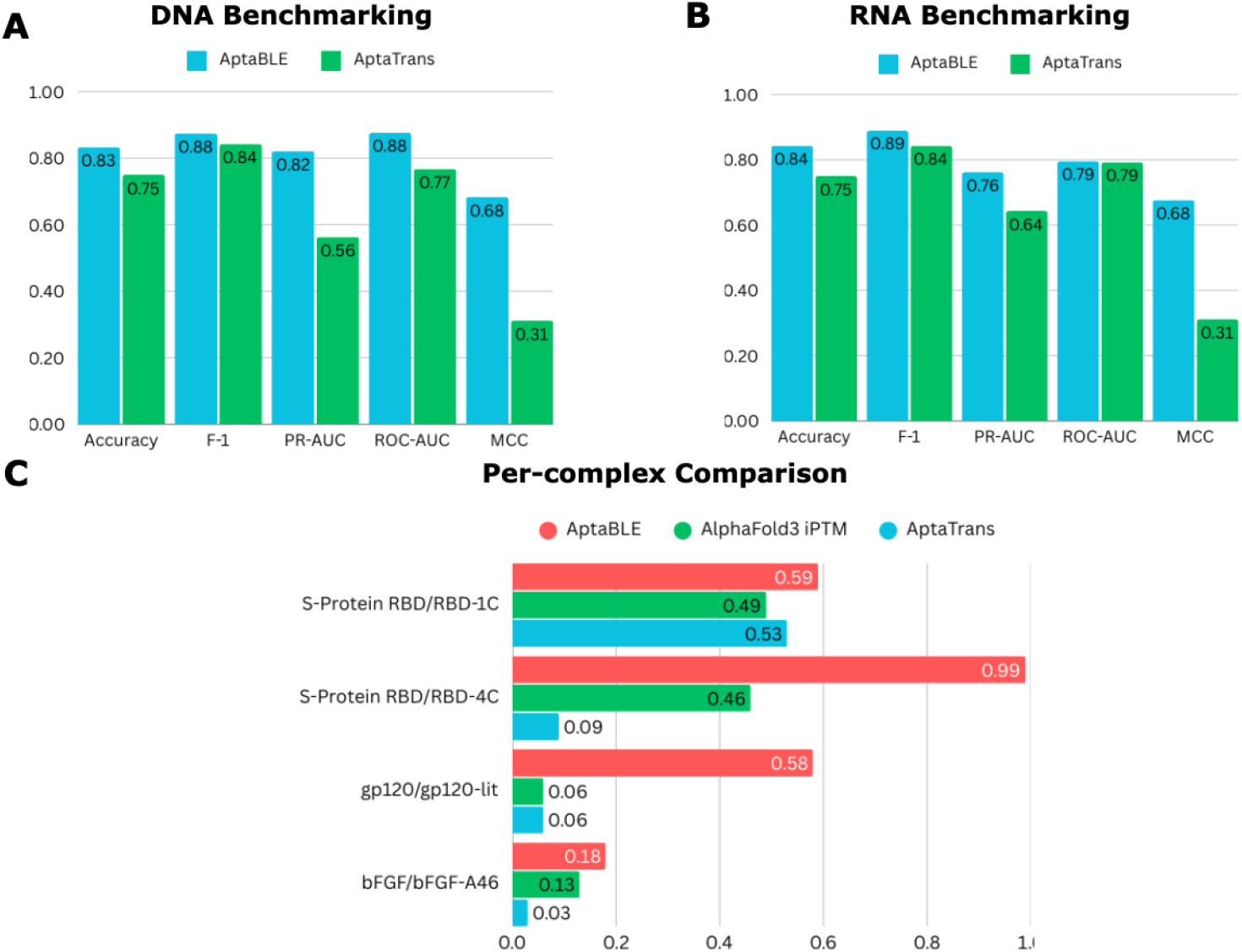
AptaBLE benchmarking. **A)** Performance of baseline models across validated binding and non-binding ssDNA aptamer datasets. **B)** Performance of baseline models across validated binding and non-binding RNA aptamer datasets. **C)** Comparison of key model metrics across four well-studied aptamer-protein systems.

**Figure S2.**
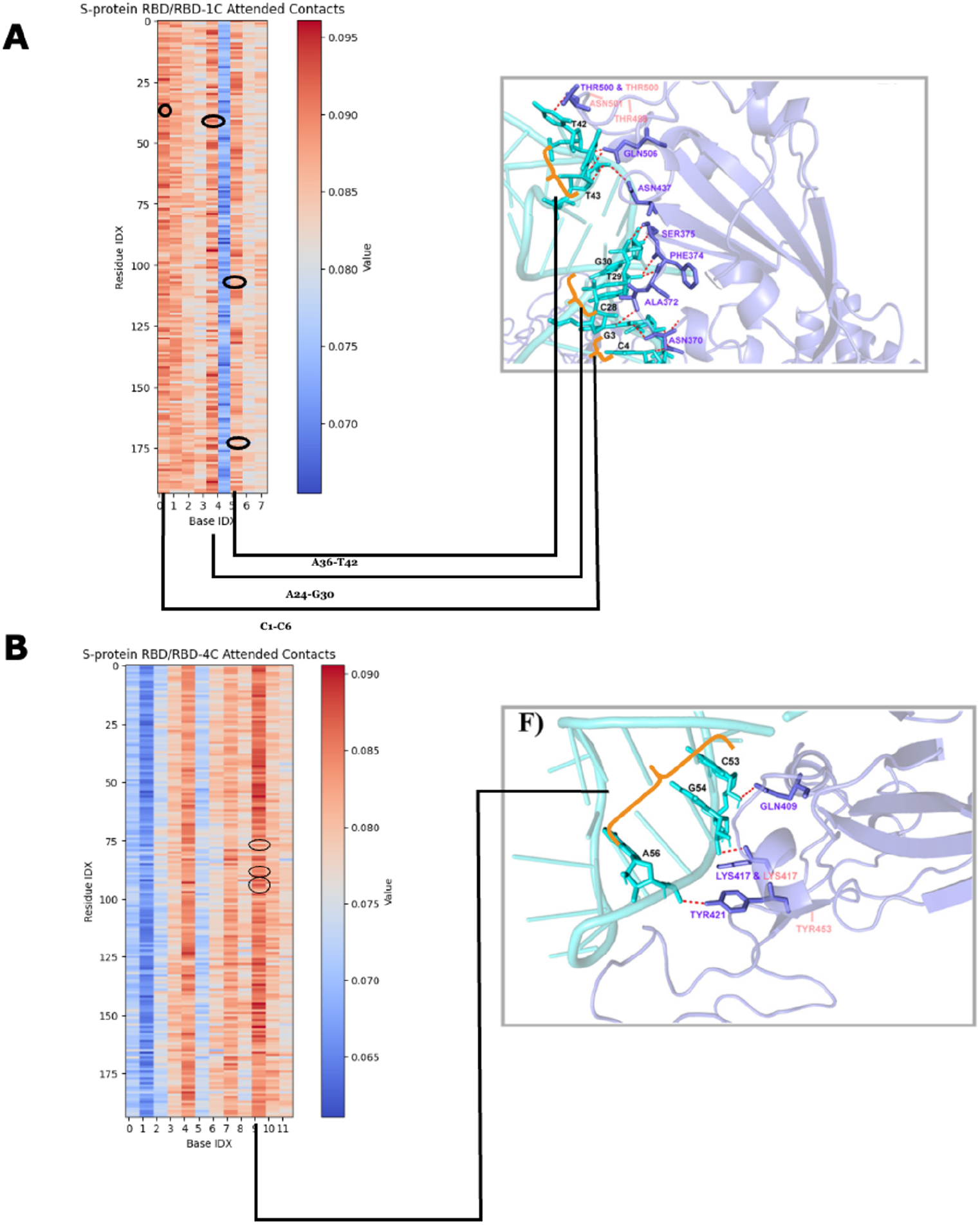
Connecting cross-attention maps to protein-aptamer binding interfaces. **A)** A cross attention map from one attention head generated from predicting RBD-1C binding to *SARS CoV-2* S-Protein RBD and the experimentally-determined binding interface. **B)** A cross attention map from one attention head generated from predicting RBD-4C binding to *SARS CoV-2* S-Protein RBD and the experimentally-determined binding interface.

**Figure S3:**
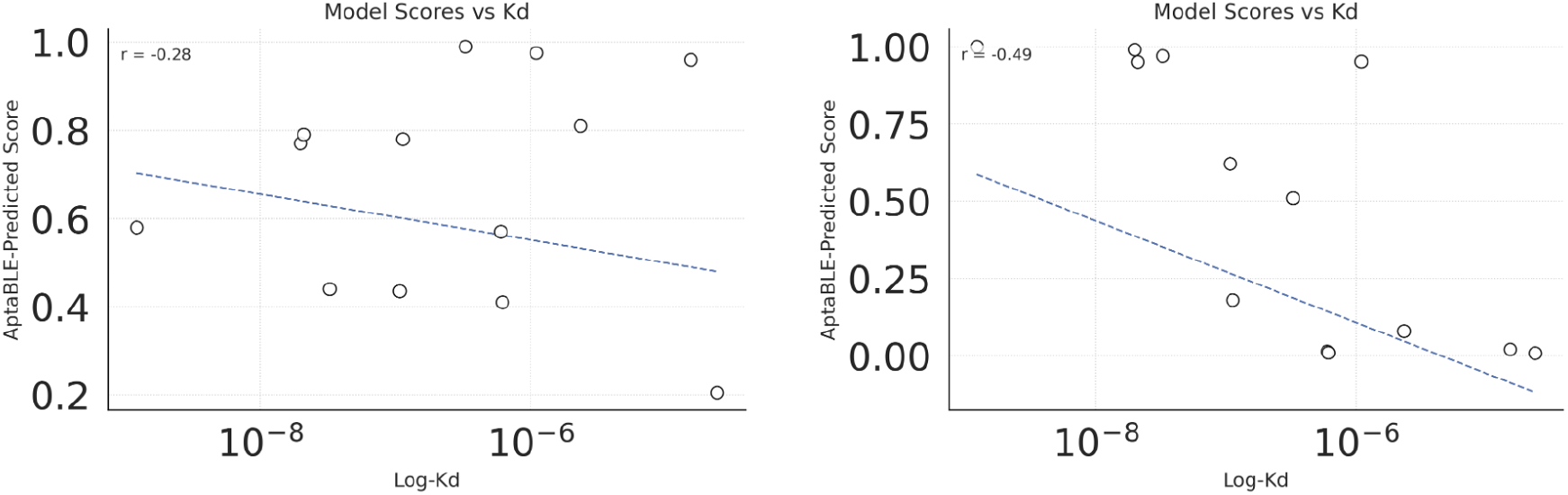
Pearson correlations for model scores and aptamer dissociation constants (*K*_*d*_). Scatterplots depict the Pearson correlation between the original (left) and fine-tuned (right) model scores and hCD117 aptamer *K*_*d*_. One aptamer originally included in **Figure 1B** was not included here as a *K*_*d*_ was not fit for it.

**Figure S4.**
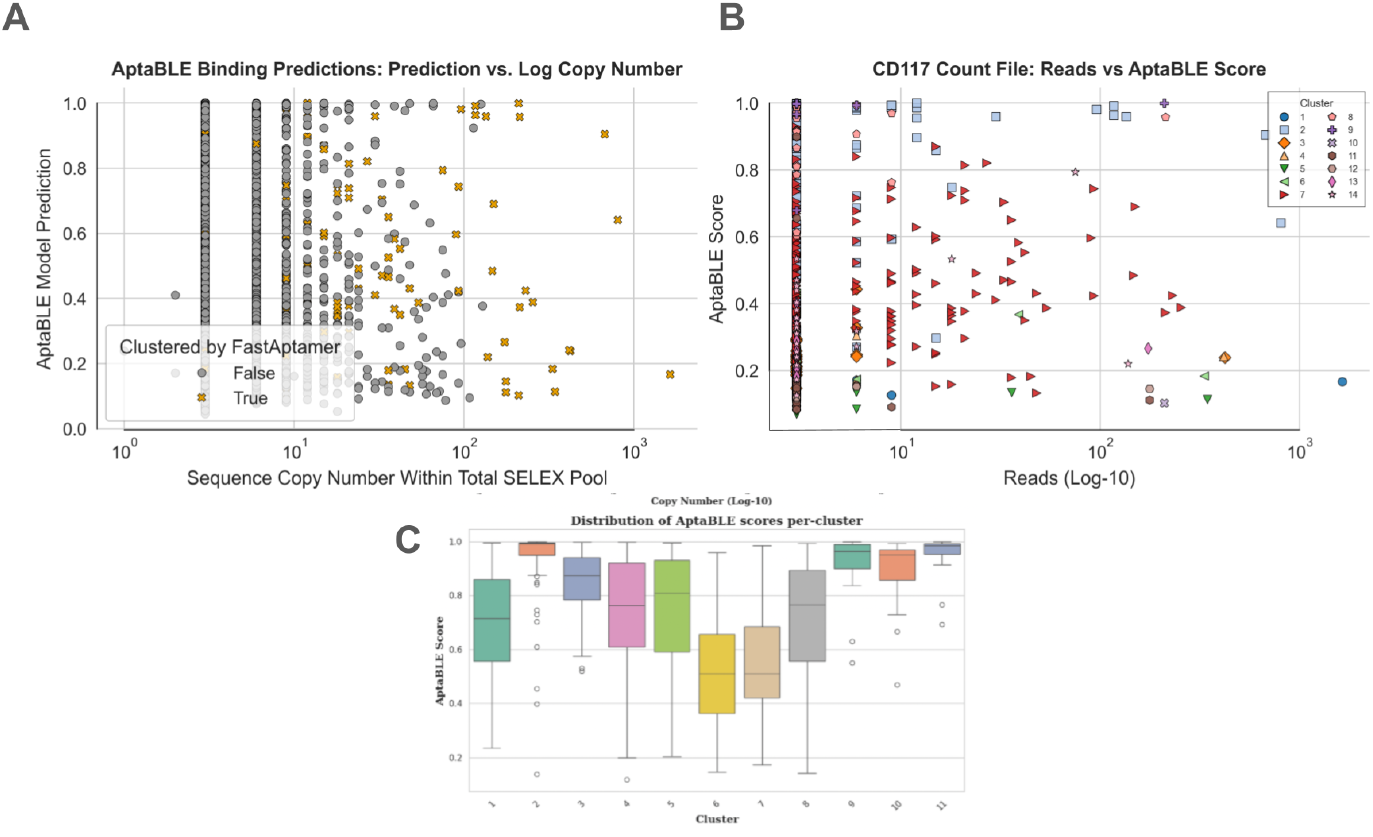
ssDNA Aptamers sequenced following nine rounds of SELEX selection for targeting hCD117. **A)** hCD117 aptamers from round 9 colored according to whether they were clustered or not. Clustering was sequence-based and performed via FastAptamer with a maximum edit distance set to 3. Red lines indicate selection thresholds used for identifying initial selection of unclustered aptamers for testing, with minimum AptaBLE score set to 0.4 and minimum copy number set to 10 reads. Encircled unclustered aptamers were chosen for binding experiments. **B)** Clustered aptamers from FastAptamer are shown, colored by their cluster. Circled aptamers were the most frequently observed within their respective cluster, according to the number of copies. **C)** AptaBLE score distributions for each FastAptamer cluster.

**Figure S5.**
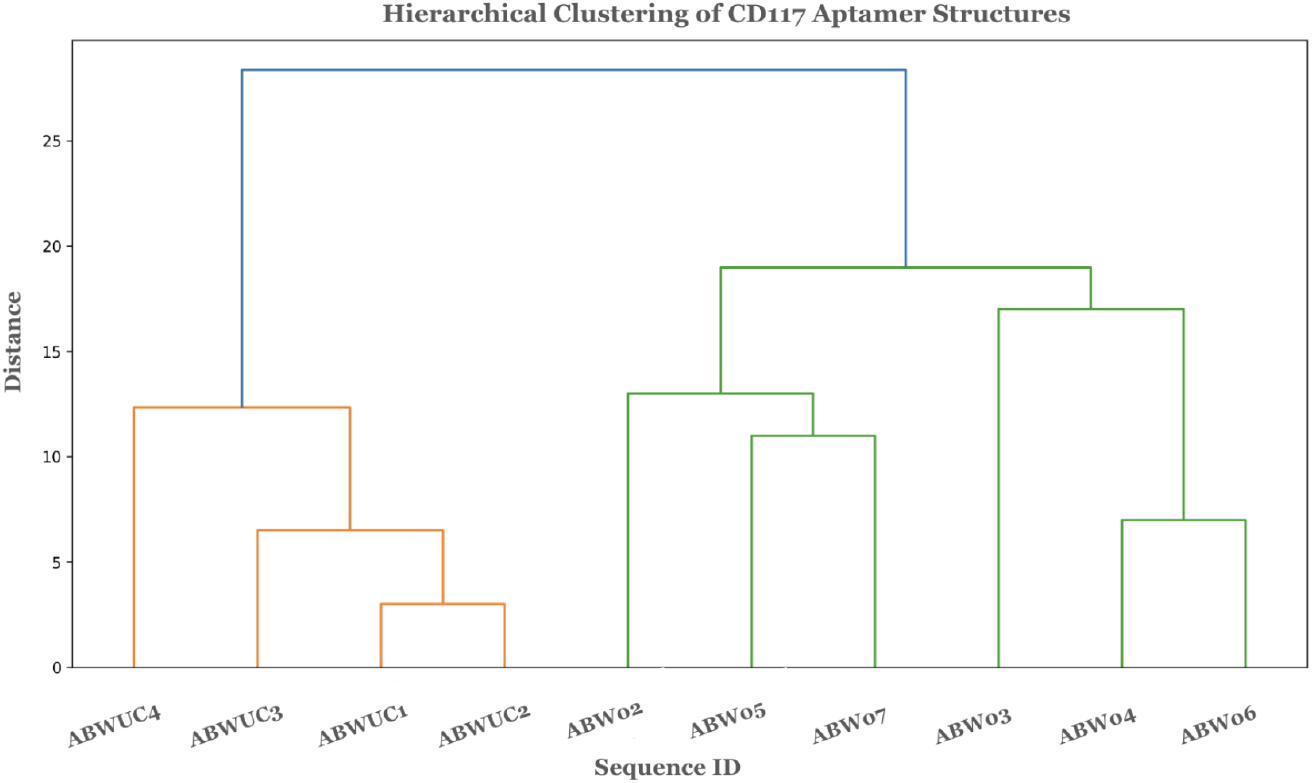
Hierarchical clustering of all tested aptamers according to their predicted secondary structures from dot-bracket representation. The 4 unclustered aptamers advanced for binding experiments are labeled as ‘ABWUC*’.

**Figure S6:**
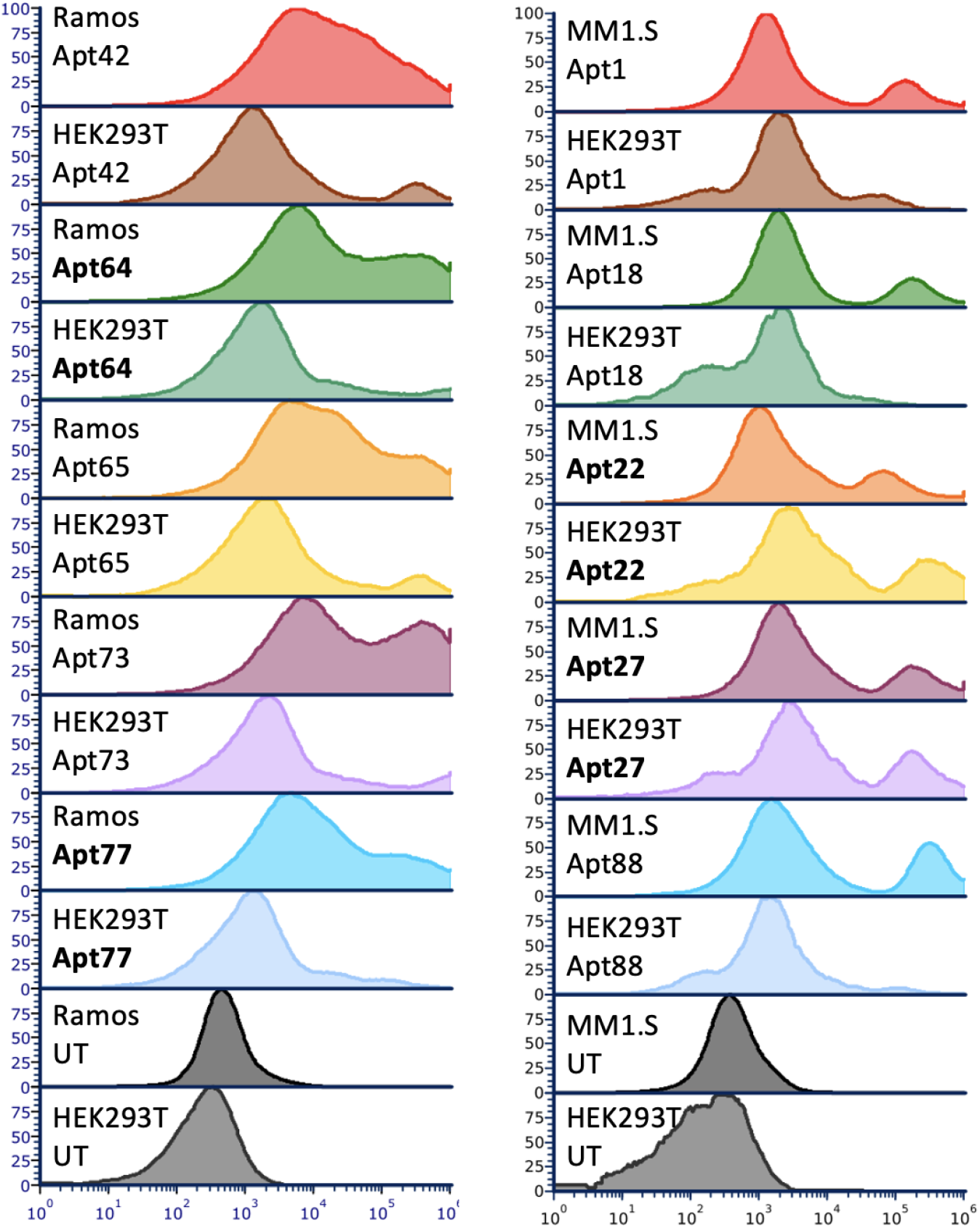
Single-concentration binding of all synthesized aptamers. Representative flow cytometry histograms show fluorescence intensity distributions for aptamers predicted to bind CD25 (**left**) and TIGIT (**right**) at 10 μM. Untreated cells (UT) serve as negative controls.

**Figure S7.**
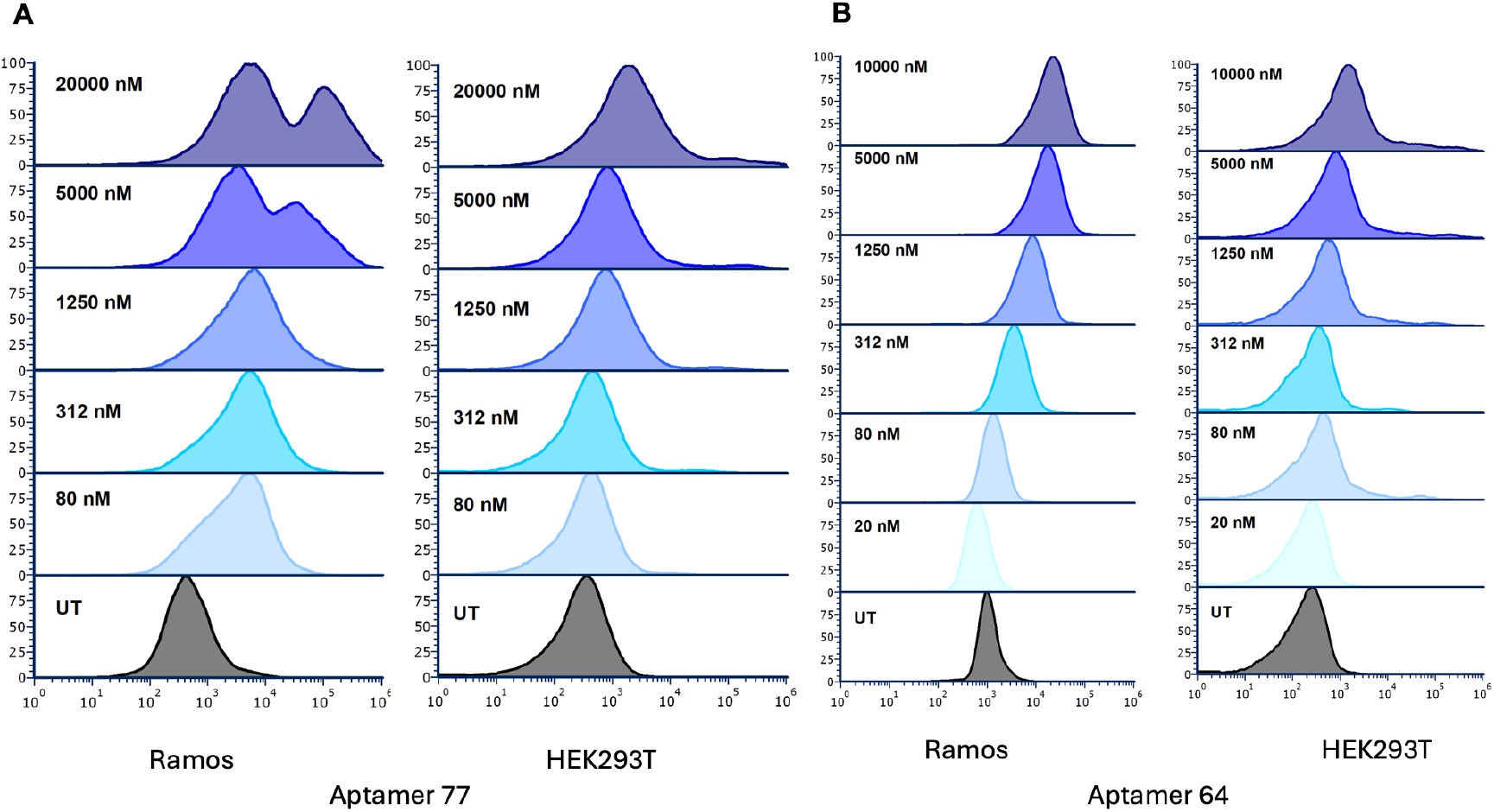
Concentration-dependent binding of Aptamers 77 and 64. Representative flow cytometry histograms showing fluorescence intensity distributions for **A)** Aptamer 77 and **B)** Aptamer 64 following incubation with Ramos (**left**) and HEK293T (**right**) cells at the indicated concentrations (80–20,000 nM). Untreated cells (UT) serve as negative controls.

**Figure S8.**
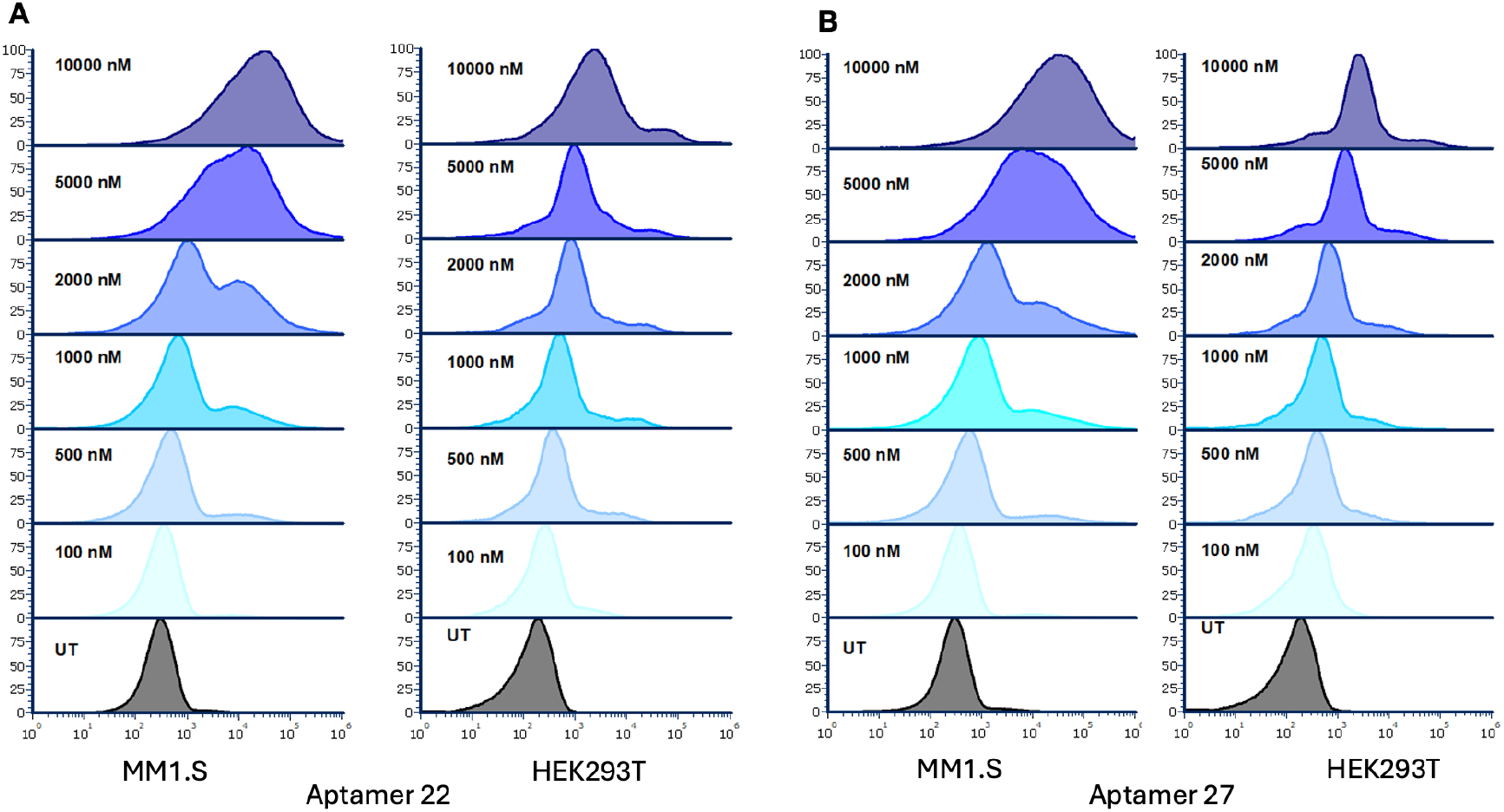
Concentration-dependent binding of Aptamers 22 and 27. Representative flow cytometry histograms showing fluorescence intensity distributions for **A)** Aptamer 22 and **B)** Aptamer 27 following incubation with MM1.S (**left**) and HEK293T (**right**) cells at the indicated concentrations (80–20,000 nM). Untreated cells (UT) serve as negative controls.

**Figure S9.**
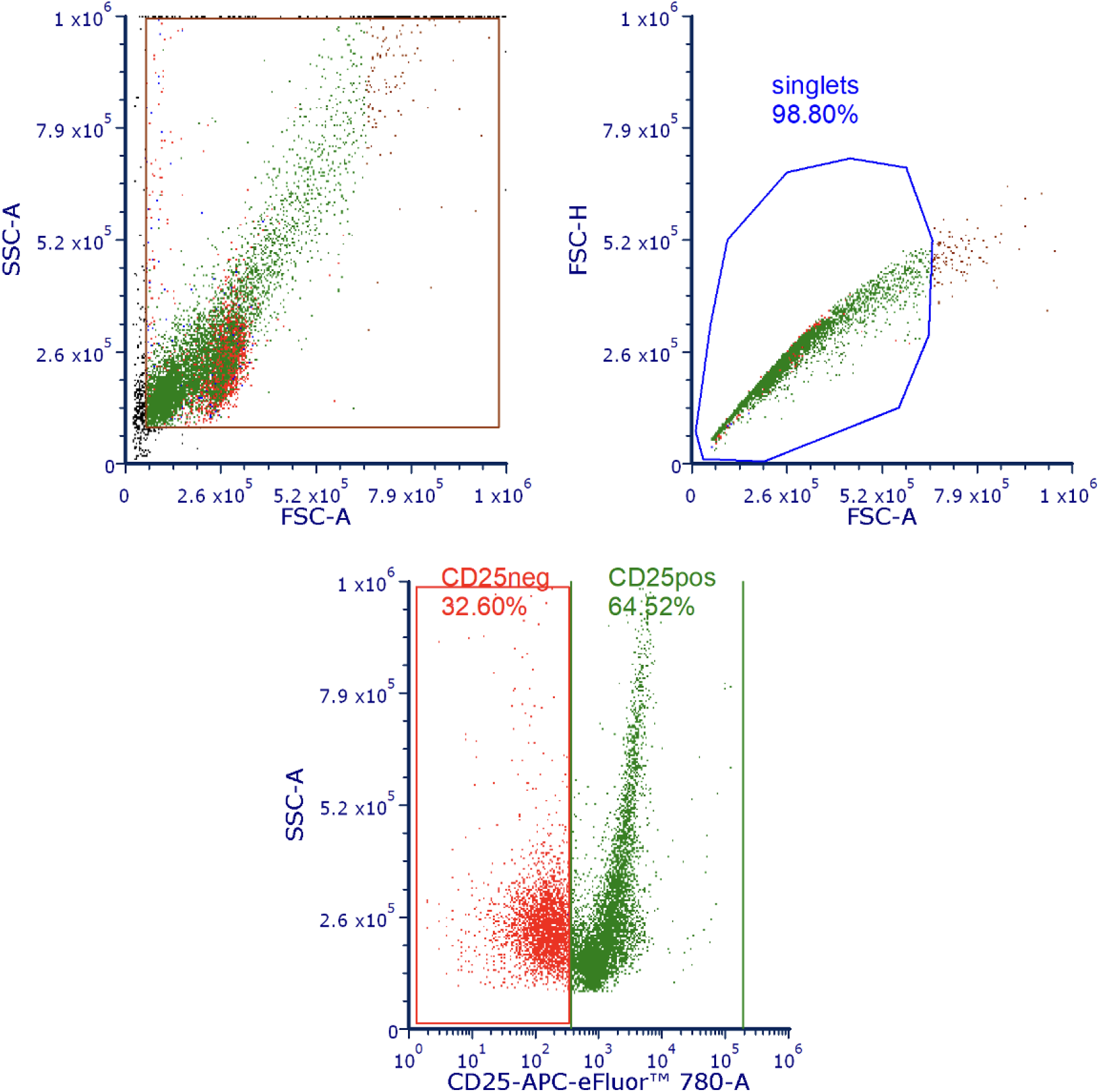
Representative gating strategy showing forward and side scatter (FSC–SSC) selection, singlet discrimination (FSC-A vs FSC-H), and CD25 expression in purified CD4^+^ T cells following treatment. CD25^+^ and CD25^−^ populations were identified based on CD25-APC-eFluor™ 780 staining to assess aptamer-mediated targeting effects.

**Figure S10.**
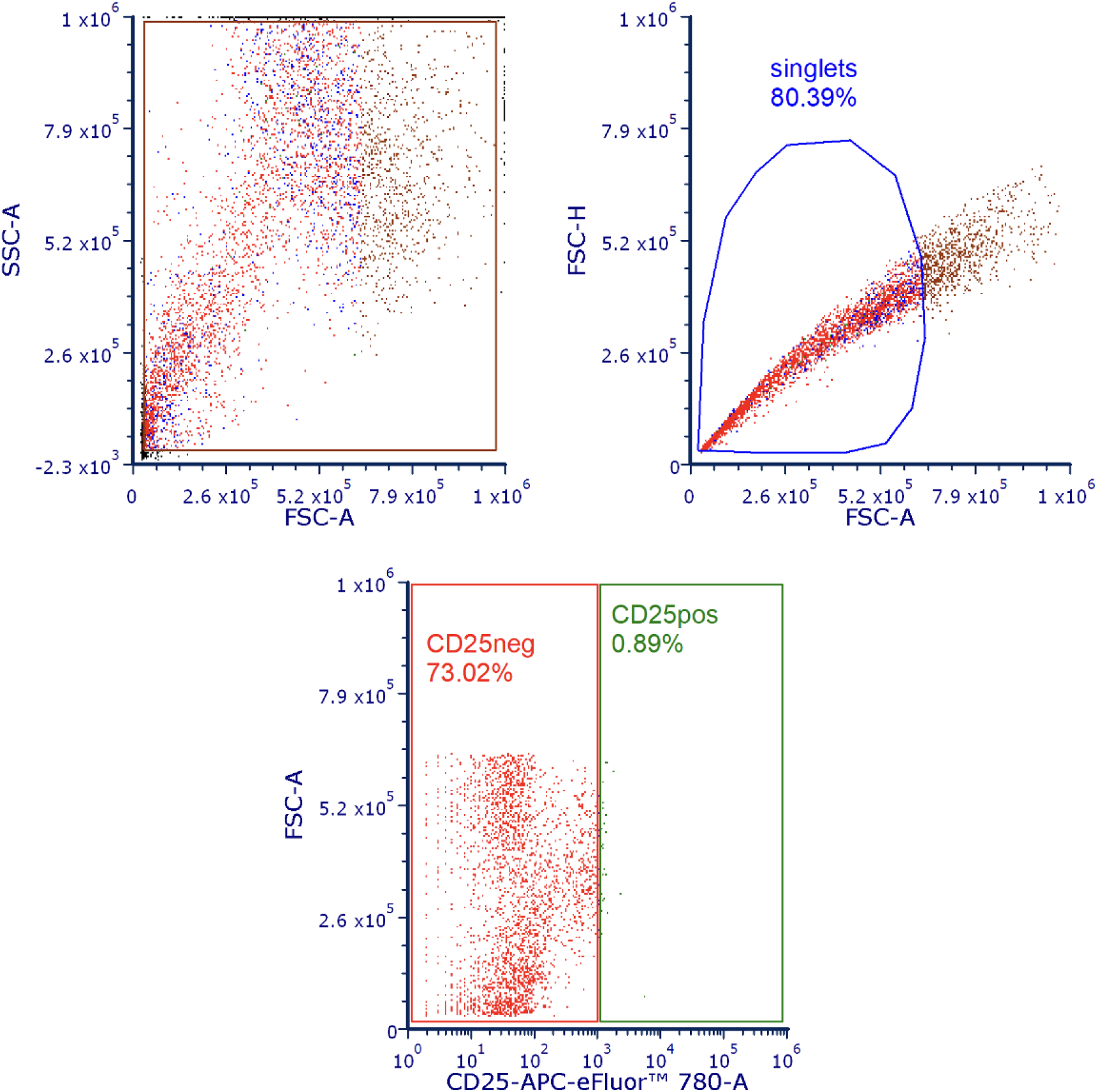
Representative FSC–SSC and singlet gating of Raw264.7 cells, followed by CD25 expression analysis. As Raw264.7 cells lack CD25 expression, these data serve as a negative control to evaluate nonspecific effects of aptamer treatment.

**Figure S11.**
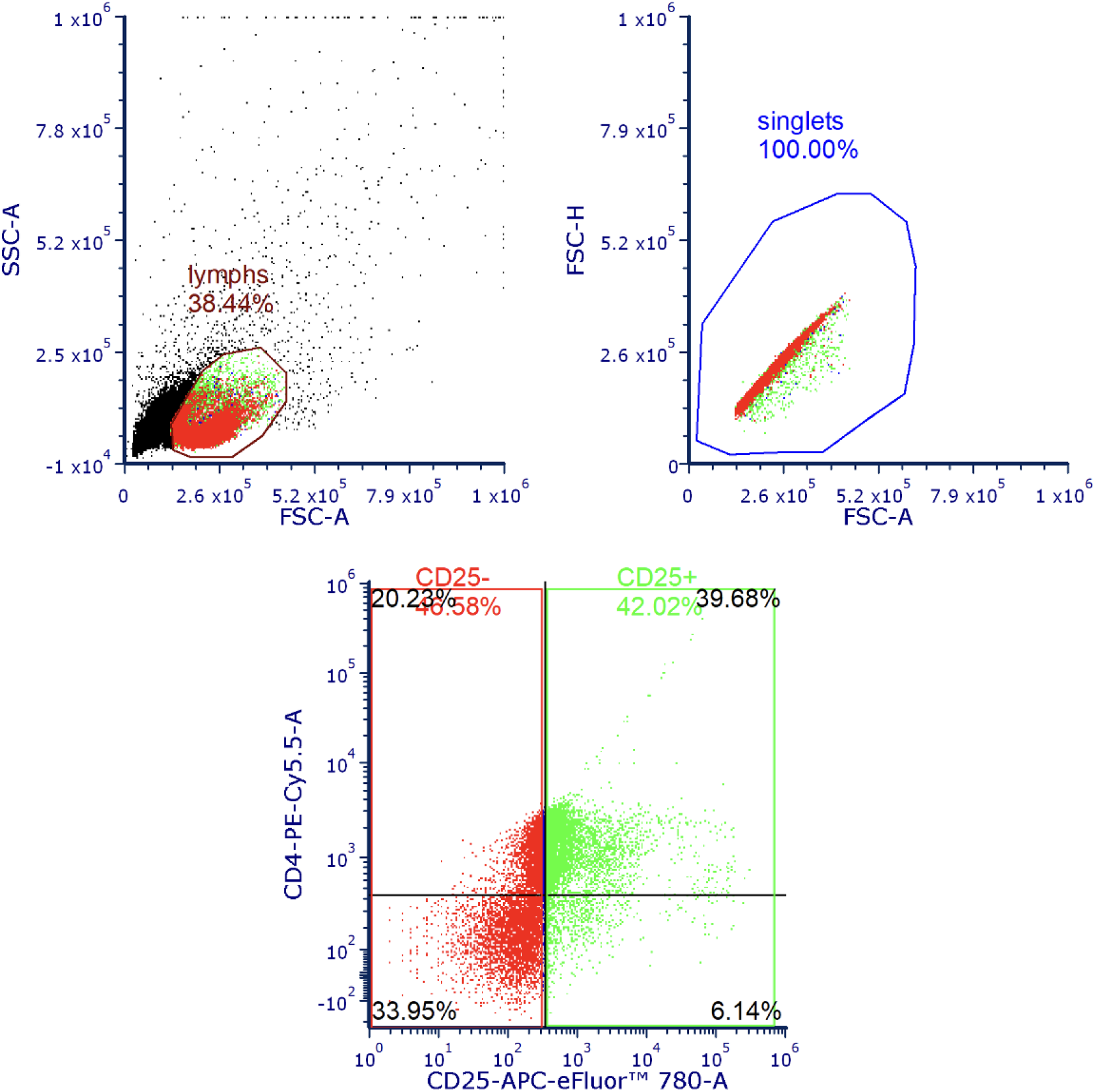
Representative gating strategy for peripheral blood mononuclear cells (PBMCs), including lymphocyte gating by FSC–SSC, singlet discrimination, and subsequent analysis of CD25 expression within CD4^+^ T-cell subsets. CD25^+^ and CD25^−^ populations were quantified to assess selective targeting in a heterogeneous immune cell population.

## References

1. Tuerk, C. & Gold, L. Systematic evolution of ligands by exponential enrichment: RNA ligands to bacteriophage T4 DNA polymerase. Science 249, 505–510 (1990).

2. Zhou, J. & Rossi, J. Aptamers as targeted therapeutics: current potential and challenges. Nat. Rev. Drug Discov. 16, 181–202 (2017).

3. Ellington, A. D. & Szostak, J. W. In vitro selection of RNA molecules that bind specific ligands. Nature 346, 818–822 (1990).

4. Thiel, W. H., Bair, T., Wyatt Thiel, K., Dassie, J. P., Rockey, W. M., Howell, C. A., Liu, X. Y., Dupuy, A. J., Huang, L., Owczarzy, R., Behlke, M. A., McNamara, J. O. & Giangrande, P. H. Nucleotide bias observed with a short SELEX RNA aptamer library. Nucleic Acid Ther. 21, 253–263 (2011).

5. Jarosch, F., Buchner, K. & Klussmann, S. In vitro selection using a dual RNA library that allows primerless selection. Nucleic Acids Res. 34, e86 (2006).

6. Hoinka, J., Berezhnoy, A., Dao, P., Sauna, Z. E., Gilboa, E. & Przytycka, T. M. Large scale analysis of the mutational landscape in HT-SELEX improves aptamer discovery. Nucleic Acids Res. 43, 5699–5707 (2015).

7. Dupuis, N. F., Holmstrom, E. D. & Nesbitt, D. J. Molecular-crowding effects on single-molecule RNA folding/unfolding thermodynamics and kinetics. Proc. Natl Acad. Sci. USA 111, 8464–8469 (2014).

8. Ishida, R., Adachi, T., Yokota, A., Yoshihara, H., Aoki, K., Nakamura, Y. & Hamada, M. RaptRanker: in silico RNA aptamer selection from HT-SELEX experiment based on local sequence and structure information. Nucleic Acids Res. 48, e82 (2020).

9. Alam, K. K., Chang, J. L. & Burke, D. H. FASTAptamer: A bioinformatic toolkit for high-throughput sequence analysis of combinatorial selections. Mol. Ther. Nucleic Acids 4, e230 (2015).

10. Wong, F., He, D., Krishnan, A., Hong, L., Wang, A. Z., Wang, J., Hu, Z., Omori, S., Li, A., Rao, J., et al. Deep generative design of RNA aptamers using structural predictions. Nat. Comput. Sci. 4, 863–872 (2024).

11. Lin, Z., Akin, H., Rao, R., Hie, B., Zhu, Z., Lu, W., Smetanin, N., dos Santos Costa, A., Fazel-Zarandi, M., Sercu, T., Candido, S. & Rives, A. Language models of protein sequences at the scale of evolution enable accurate structure prediction. bioRxiv (2022). doi:10.1101/2022.07.20.500902

12. Dalla-Torre, H., Gonzalez, L., Mendoza-Revilla, J., Lopez Carranza, N., Grywaczewski, A. H., Oteri, F., Dallago, C., Trop, E., Sirelkhatim, H., Richard, G., et al. The nucleotide transformer: Building and evaluating robust foundation models for human genomics. bioRxiv (2023). doi:10.1101/2023.01.11.523679

13. Chen, J., Hu, Z., Sun, S., Tan, Q., Wang, Y., Yu, Q., Zong, L., Hong, L., Xiao, J., King, I. & Zhang, Y. Interpretable RNA foundation model from unannotated data for highly accurate RNA structure and function predictions. arXiv preprint arXiv:2204.00300 (2022).

14. Shin, I., Kang, K., Kim, J., Sel, S., Choi, J., Lee, J.-W., Kang, H. Y. & Song, G. AptaTrans: a deep neural network for predicting aptamer-protein interaction using pretrained encoders. BMC Bioinformatics 24, 447 (2023).

15. Woldeyes, R. A., Sivak, D. A. & Fraser, J. S. E pluribus unum, no more: from one crystal, many conformations. Curr. Opin. Struct. Biol. 28, 56–62 (2014).

16. Hermann, T. & Patel, D. J. Adaptive recognition by nucleic acid aptamers. Science 287, 820–825 (2000).

17. Corso, G., Stärk, H., Jing, B., Barzilay, R. & Jaakkola, T. DiffDock: Diffusion steps, twists, and turns for molecular docking. arXiv preprint arXiv:2210.01776 (2022).

18. Lu, W., Wu, Q., Zhang, J., Rao, J., Li, C. & Zheng, S. TANKBind: Trigonometry-aware neural networks for drug-protein binding structure prediction. Adv. Neural Inf. Process. Syst. 35, 7236–7249 (2022).

19. Cao, D., Chen, G., Jiang, J., Yu, J., Zhang, R., Chen, M., Zhang, W., Chen, L., Zhong, F., Zhang, Y., et al. Generic protein-ligand interaction scoring by integrating physical prior knowledge and data augmentation modelling. Nat. Mach. Intell. 6, 1439–1453 (2024).

20. Abramson, J., Adler, J., Dunger, J., Evans, R., Green, T., Pritzel, A., Ronneberger, O., Willmore, L., Ballard, A. J., Bambrick, J., et al. Accurate structure prediction of biomolecular interactions with AlphaFold 3. Nature 630, 493–500 (2024).

21. Jumper, J., Evans, R., Pritzel, A., Green, T., Figurnov, M., Ronneberger, O., Tunyasuvunakool, K., Bates, R., Žídek, A., Potapenko, A., et al. Highly accurate protein structure prediction with AlphaFold. Nature 596, 583–589 (2021).

22. Dwivedy, A., Baskaran, D., Sharma, G., Hong, W., Gandavadi, D., Krissanaprasit, A., … & Wang, X. (2025). Engineering Novel DNA Nanoarchitectures for Targeted Drug Delivery and Aptamer Mediated Apoptosis in Cancer Therapeutics. Advanced Functional Materials, 35(22), 2425394.

23. Lee, G., Jang, G. H., Kang, H. Y. & Song, G. Predicting aptamer sequences that interact with target proteins using an aptamer-protein interaction classifier and a Monte Carlo tree search approach. PLoS One 16, e0253760 (2021).

24. Sophia Tang, Yinuo Zhang, and Pranam Chatterjee. Peptune: De novo generation of therapeutic peptides with multi-objective-guided discrete diffusion. ArXiv, 2412, (2025).

25. Xu, M., Yuan, X., Miret, S. & Tang, J. ProtST: Multi-modality learning of protein sequences and biomedical texts. Proc. 40th Int. Conf. Mach. Learn. 202, 38749–38767 (2023).

26. Patel, S., Peng, F. Z., Fraser, K., Friedman, A. D., Chatterjee, P., & Yao, S. (2025). EvoFlow-RNA: Generating and Representing non-coding RNA with a Language Model. bioRxiv, 2025–02.

27. Askari, A., Kota, S., Ferrell, H., Swamy, S., Goodman, K. S., Okoro, C. C., Crenshaw, I. C. S., Hernandez, D. K., Oliphant, T. E., Badrayani, A. A., et al. UTExas Aptamer Database: the collection and long-term preservation of aptamer sequence information. Nucleic Acids Res. 52, D351–D359 (2024).

28. Li, B.-Q., Zhang, Y.-C., Huang, G.-H., Cui, W.-R., Zhang, N. & Cai, Y.-D. Prediction of aptamer-target interacting pairs with pseudo-amino acid composition. PLoS One 9, e86729 (2014).

29. Lorenz, R., Bernhart, S. H., Höner zu Siederdissen, C., Tafer, H., Flamm, C., Stadler, P. F. & Hofacker, I. L. ViennaRNA Package 2.0. Algorithms Mol. Biol. 6, 26 (2011).

30. Steinegger, M. & Söding, J. MMseqs2 enables sensitive protein sequence searching for the analysis of massive data sets. Nat. Biotechnol. 35, 1026–1028 (2017).

31. Lin, Y.-C., Chen, W.-Y., Hwu, E.-T. & Hu, W.-P. In-silico selection of aptamer targeting SARS-CoV-2 spike protein. Int. J. Mol. Sci. 23, 5810 (2022).

32. Zhao, N., Pei, S. N., Parekh, P., Salazar, E., & Zu, Y. (2014). Blocking interaction of viral gp120 and CD4-expressing T cells by single-stranded DNA aptamers. The international journal of biochemistry & cell biology, 51, 10–18.

33. Eguchi, A., Ueki, A., Hoshiyama, J., Kuwata, K., Chikaoka, Y., Kawamura, T., … & Sando, S. (2021). A DNA aptamer that inhibits the aberrant signaling of fibroblast growth factor receptor in cancer cells. Jacs Au, 1(5), 578–585.

34. Giulini, M., Reys, V., Teixeira, J. M., Jiménez-García, B., V. Honorato, R., Kravchenko, A., … & Bonvin, M. (2025). HADDOCK3: A modular and versatile platform for integrative modeling of biomolecular complexes. Journal of Chemical Information and Modeling.

35. Pettersen, E. F., Goddard, T. D., Huang, C. C., Meng, E. C., Couch, G. S., Croll, T. I., … & Ferrin, T. E. (2021). UCSF ChimeraX: Structure visualization for researchers, educators, and developers. Protein science, 30(1), 70–82.

